# Stepwise remodelling and subcompartment formation in individual vesicles by three ESCRT-III proteins

**DOI:** 10.1101/2022.02.25.481928

**Authors:** Yunuen Avalos-Padilla, Vasil N. Georgiev, Eleanor Ewins, Tom Robinson, Esther Orozco, Reinhard Lipowsky, Rumiana Dimova

**Affiliations:** Department of Theory and Bio-Systems, Max Planck Institute of Colloids and Interfaces, Science Park Golm, 14476 Potsdam, Germany; Nanomalaria Group, Institute for Bioengineering of Catalonia (IBEC), The Barcelona Institute of Science and Technology, Baldiri Reixac 10-12, ES-08028 Barcelona, Spain; Barcelona Institute for Global Health (ISGlobal, Hospital Clínic-Universitat de Barcelona), Rosselló 149-153, ES-08036 Barcelona, Spain; Departamento de Infectómica y Patogénesis Molecular, CINVESTAV IPN, Ciudad de México, México

**Keywords:** GUV, intraluminal vesicles, bending rigidity, membrane tension, protein domain, microfluidics, micropipette aspiration

## Abstract

The endosomal sorting complex required for transport (ESCRT) is a multi-protein complex involved in several membrane remodelling processes. Different approaches have been used to dissect the mechanism by which ESCRT proteins produce scission in the membranes. However, the underlying mechanisms generating the membrane deformations remain poorly understood. In this study, giant unilamellar vesicles (GUVs), microfluidic technology and micropipette aspiration are combined to continuously follow the ESCRT-III-mediated membrane remodelling on the single-vesicle level for the first time. With this approach, we identify different mechanisms by which a minimal set of three ESCRT-III proteins from the phagocytic parasite *Entamoeba histolytica* reshape the membrane. These proteins modulate the membrane stiffness and spontaneous curvature to regulate the bud size and generate intraluminal vesicles in GUVs even in the absence of ATP. We show that the bud stability depends on the protein concentration and membrane tension. The approach introduced here should open the road to diverse applications in synthetic biology for establishing artificial cells with several membrane compartments.

## 1. Introduction

The endosomal sorting complex required for transport (ESCRT) was first described as a vacuolar protein sorting machinery in yeast ^[1]^. Over the years, it has been demonstrated that the ESCRT machinery participates during membrane fission and remodelling in different processes including the formation of multivesicular bodies ^[2]^, virus budding ^[3]^, neuron pruning ^[4]^, plasma membrane repair ^[5]^, and autophagy ^[6]^ as reviewed in ^[7]^. The ESCRT machinery is a multi-protein system formed by the sub-complexes ESCRT-0, ESCRT-I, ESCRT-II, ESCRT-III and a set of accessory proteins that act in a sequential manner to generate invaginations in the membrane^[8]^. Among these sub-complexes, ESCRT-III is the only protein family that remodels the membrane shape and it is also the most conserved across the eukaryotic lineage. Moreover, some members of the ESCRT-III family have been found in the Archaea taxa suggesting its ancestral function as a scission machinery ^[9]^. The ESCRT-III complex is formed by Snf7-domain-containing proteins and the number of components varies among the different supergroups within the eukaryotic taxa, probably due to specialized evolution in the diverse organisms. The most essential proteins of ESCRT-III are Vps2, Vps20, Vps24 and Snf7/Vps32. All of them share a common structural core motif formed by at least four alpha helices with a positive net charge (core domain) with the propensity to bind to negatively charged lipid membranes ^[10]^. ESCRT-III proteins also possess a negatively charged fifth alpha helix that blocks the core domain thus, allowing the proteins to remain soluble in the cytoplasm in the absence of activating factors ^[11]^.

Although membrane fission conducted by ESCRT-III is not fully understood, it is generally believed that ESCRT-III polymers bind transiently to highly curved regions of membranes ^[12]^, and grow toward zones with less curvature ^[13]^. The constriction of the bud neck is mediated by the formation of Vps32 polymers, later remodelled by Vps24, Vps2 ^[14]^ and the ATPase Vps4 ^[15]^ to produce domes and cones ^[16]^. Theoretical estimates suggest that the formation of cones is more favoured since it requires less adhesive strength of the protein-membrane interaction and is associated with higher constriction forces ^[17]^.

Giant unilamellar vesicles (GUVs) ^[18]^ represent a well-established tool not only for elucidating membrane properties and remodelling ^[18b,19]^, but also as biomimetic containers in synthetic biology ^[20]^. Different studies using GUVs and the purified components of the ESCRT machinery have suggested that ESCRT-III is able to induce invaginations in membranes without the requirement of the upstream ESCRT factors ^[21]^. Additionally, we have recently shown that the core domain of the recombinant Vps20 from *Entamoeba histolytica*, EhVps20(1-173) (hereafter referred to as EhVps20t), can bind to the membrane of GUVs, and together with EhVps32 and EhVps24 they are sufficient to generate vesicular subcompartments (which we will refer to as intraluminal vesicles, ILVs) in the same GUV ^[21c]^. Moreover, by using the same system, we have observed that alterations to the order of protein addition showed no significant differences but omission of any of them resulted in no ILV formation^[21c]^. Despite these findings, the factors governing the size of the intraluminal vesicles and the role of the membrane material properties in regulating the ESCRT-III activity are unknown. Indeed, knowing these factors should be useful in order to construct cell-size vesicles with nested compartments for the reconstitution of the structural mimicry of eukaryotes and their membrane-bound organelles. Such synthetic compartmentalization offers a route towards uncoupling enzymatic reactions or separating reagents. While efforts in this direction have been already made, see e.g. ^[22]^, the most successful case, allowing for control on the compartment size and number, relies on the implementation of microfluidics on double emulsions for the preparation of vesicles-in-vesicle (vesosome) systems ^[22a]^. The drawback of this approach, in which the vesicles are constructed in a layer-by-layer fashion, has certain disadvantages for protein reconstitution. We speculate that closer-to-nature generation of internal micron-sized compartments as those triggered by ESCRTs and regulating their properties via modulating determinants such as membrane composition and rigidity will pave the road towards more natural routes for creating synthetic cells with multiple compartments. In the present study, we combined cellular and synthetic biology approaches to elucidate the mechanism for membrane budding and fission triggered by a minimal set of three ESCRT-III proteins from the highly phagocytic parasite *E. histolytica*. Using a single-vesicle assay, we identify three main steps of the membrane reshaping process: the first ESCRT-III component binds to the membrane, the second component binds to the first one and generates inward pointing buds, and the third one leads to membrane fission even in the absence of ATP (suggesting that cells can take advantage of passive processes). We show that the size of the generated ILVs does not depend exclusively on the size of the engulfed cargo, as previously suggested ^[21a]^, but is also influenced by the membrane mechanical properties and the protein coverage. The stability and reversible formation of intraluminal invaginations was probed against increased membrane tension employing osmotic inflation/deflation and micropipette aspiration of giant vesicles. The system described here offers a minimalistic approach for establishing a synthetic microcompartmentalized cell, in which the size and content of the compartments can be externally controlled.

## 2. Results

### 2.1. Fission triggered by ESCRT-III proteins can be divided in three consecutive steps: protein binding, membrane budding and fission

Previous studies^[21b, 21c, 23]^ have investigated the simultaneous action of protein mixtures on GUV morphology, thereby probing the membrane response averaged over a batch of vesicles. This averaging procedure does not allow several important aspects of the membrane remodelling process to be addressed: Did the addition of one of the proteins compromise the membranes in terms of vesicle leakage and permeation? What was the initial morphology of the GUVs before they started to interact with the proteins? Indeed, vesicles in the same batch also have different membrane tensions typically in the range between 10^-9^ and 10^-4^ N/m and exhibit various morphologies depending on the leaflet asymmetry across the bilayer membrane ^[24]^ (they may even vary in composition when the membrane contains several molecular components ^[25]^). Thus, we investigated the action of proteins on the same individual vesicle by adding them consecutively to this vesicle. To avoid possible effects of unbound proteins, a microfluidic device, which allows capture of GUVs and exchange of their external solution, was used to follow the interactions on individual vesicles ^[26]^. GUVs composed of POPC:POPS:Chol:PI(3)P (62:10:25:3), roughly mimicking the endosomal membrane composition ^[27]^ and labelled with TR-DHPE were electroformed and captured between the posts in a microfluidic device; for details, see Section S1 and Figure S1 in the supporting information (SI). Thereafter, the three ESCRT-III components EhVps20t, EhVps32 and EhVps24 were sequentially flushed in at a constant flow rate of 0.1 μl/min. A total of 100 μl solution was used to obtain complete exchange at each solution-exchange step to ensure concentration control. After introducing each protein, the solution in the chamber was exchanged with the protein-free buffer (25 mM Tris, 150 mM NaCl, pH = 7.4) to remove the unbound proteins in the surrounding solution. This step aimed at resolving curvature effects specific to the particular protein added. Fluorescent analogues of EhVps20t and EhVps24 were used to monitor protein binding. Consistent with previous bulk studies on vesicle populations ^[21c]^, EhVps20t was observed to bind to the membrane of the GUVs. Even after a thorough washing, EhVps20t remains bound to the vesicle (Figure 1, first two rows of images). Note that close contact to the PDMS posts decreases the fluorescence signal from the membrane as previously observed ^[28]^. After the addition of EhVps32 and slight deflation of less than 5% to allow for excess area, small invaginations (intraluminal buds) attached to the membrane of GUVs with relatively uniform size were observed (Figure 1, third row, Movie S1 in the SI); similar deflation in the presence of EhVps20t was not found to result in detectable morphological changes in the GUVs (Figure 1, second row). Finally, the addition of ATP-free solution of EhVps24 triggered the scission of the newly formed ILVs and their release in the vesicle interior (Figure 1, bottom row). It was difficult to detect the ILVs from bright-field observations during the experiment due to spinning of the vesicles in the microfluidic flow. However, the generated ILVs could be observed upon refocusing and stopping the flow (Movie S2 in the SI). In this way, the ILVs were found to have a relatively homogeneous size distribution (1 ± 0.18 μm diameter, obtained by three independent replicas where 2 vesicles were followed) and were similar in diameter to the intraluminal buds generated after introducing EhVps32. Unlike ILVs, the size of intraluminal buds could not be measured systematically because of imaging artefacts arising from the proximity of the mother vesicle membrane but we were able to estimate their sizes from those favorable cases for which the buds are clearly visible. Incubation of GUVs with six rounds of alternating buffer flushing and incubation, resulted in no ILV formation (Figure S2), indicating that the effect is due to protein activity and not because of buffer or flow in the microfluidic chamber.

**Figure 1.**
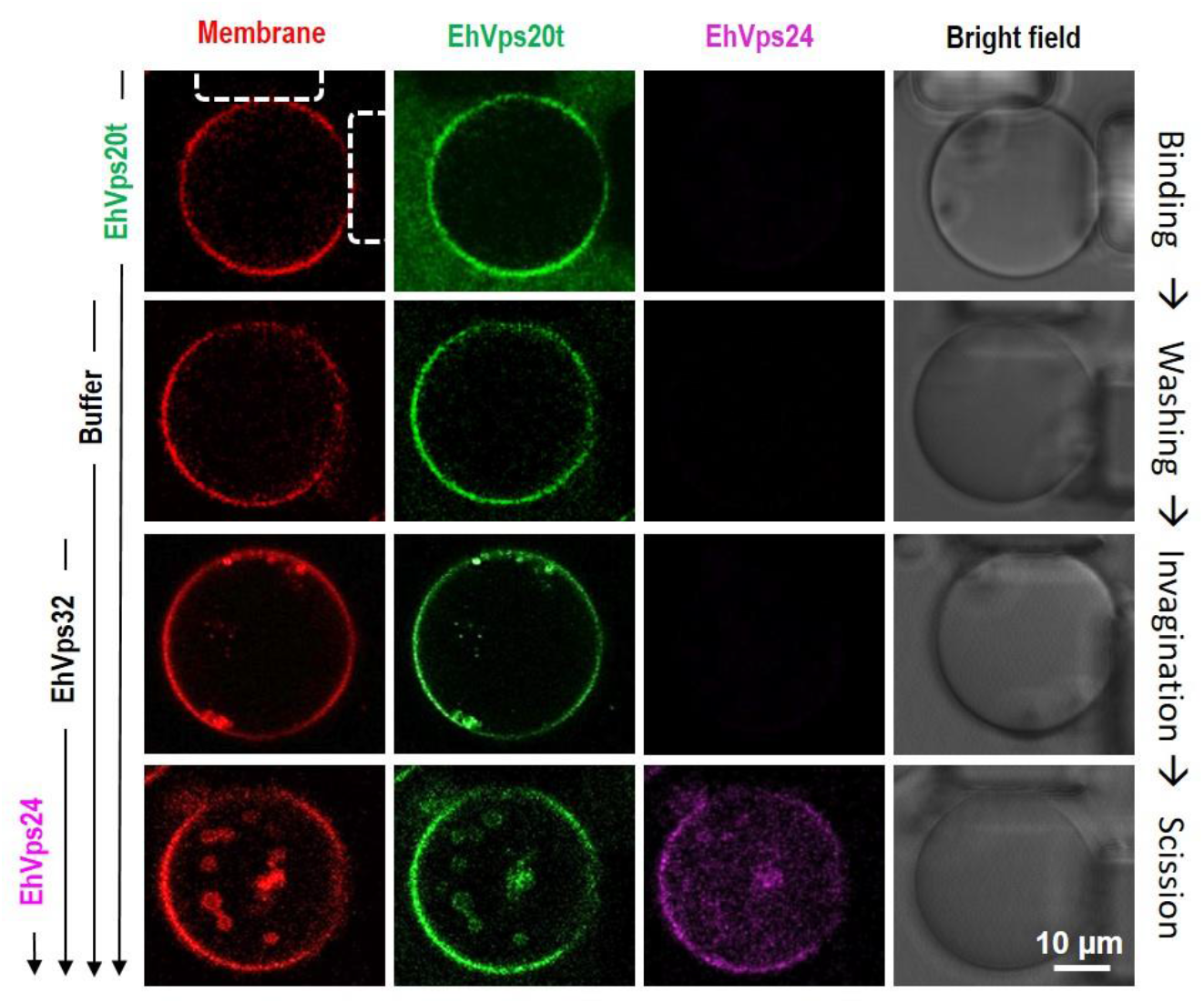
Single-vesicle assay for imaging the successive binding of ESCRT-III proteins, membrane (inward) budding and scission. Vesicles prepared from POPC:POPS:Chol:PI(3)P (62:10:25:3) and labelled with 0.1 mol% TR-DHPE were loaded on a microfluidic chamber at a flow rate of 10 μl/min. Then, 125 nM of EhVps20t (20% of the protein was labelled with Oregon green), 600 nM EhVps32 and 300 nM EhVps24 (20% of the protein was labelled with Alexa 633) were successively flushed in at a constant rate of 0.1 μl/min. Before introducing the next protein solution, the chamber was flushed with 100 μl of protein-free buffer to remove unbound proteins in the bulk as illustrated for EhVps20t with the first two rows of images. The snapshots show confocal cross sections and phase-contrast images (last column) of the same trapped vesicle after the flushing steps. Three main events are distinguished: EhVps20t binding of the membrane (first two rows), budding triggered by EhVps32 (third row) and scission directed by EhVps24 (last row). To avoid cross talk, the confocal images were obtained from sequential scanning. The microposts trapping the vesicle are seen in the last column of images and are indicated by dashed lines in the upper left image. Experiments were conducted three times and, in each experiment, at least two different vesicles were monitored for the whole sequence of flushing/incubation steps (see also Movie S1).

### 2.2. Quantifying the amount of EhVps20t bound to the membrane: dependence on protein bulk concentration

As shown previously, the interactions between ESCRT-III proteins and membranes are mostly mediated by electrostatic forces between positive residues and negatively-charged lipids ^[1c, 10b, 16a]^. Despite the selective binding of upstream ESCRT factors to the endosomal enriched lipid PI(3)P ^[29]^, several studies demonstrated that ESCRT-III proteins are able to bind to other negatively-charged lipids. In particular, the binding and function of ESCRT-III proteins is not significantly different when using membrane compositions of PC:PS:Chol:PI(3)P and PC:PS with the same negative surface charge density ^[6a, 21c, 30]^, as recently confirmed for *E. histolytica* ESCRT-III proteins^[31]^. For quantifying the amount of EhVps20t on the membrane, we used GUVs with a simpler lipid composition, namely POPC:POPS (80:20), which have a similar surface charge density as vesicle membranes made of POPC:POPS:Chol:PI(3)P (62:10:25:3); see SI Section S2.

To quantify the EhVps20t coverage, we analysed the fluorescence signal from the labelled protein and extrapolated the concentration from a calibration curve obtained for a lipid labelled with the same fluorophore, namely the Oregon green 488 labelled lipid OG-DHPE. The fluorescence signal in the membrane, *I*, measured at different pre-set molar fractions of OG-DHPE in the vesicles (**Figure 2**a, see SI Section S2 and Figures S3-S5) was found to follow the linear dependence: *I* ≅ 115 *n_OG_* – 7 [a.u.], where *n_OG_* is the mole fraction (in %) of OG label in the membrane. Then, the intensity signal from protein labelled with the same fluorescent group, OG-EhVps20t, and bound to the outer leaflet of the vesicle membrane was measured and the background signal from free protein in the bulk subtracted (see Figure S5). The molar fraction of the protein at the membrane of vesicles made of POPC:POPS (80:20) was determined using the calibration curve (taking into account that the fluorescence from the protein at the outer vesicle leaflet should be compared to half the intensity of OG-DHPE located in both leaflets). The protein coverage on vesicles prepared from POPC:POPS:Chol:PI(3)P (62:10:25:3) was found to be similar as expected from the comparable surface charge, see supplementary Figure S6. The results in Figure 2b show a Langmuir-type adsorption isotherm where the membrane coverage of EhVps20t increased with protein bulk concentration and reached saturation at around 800 nM (Figure 2b) protein in the bulk. From the obtained membrane coverage of the protein we could roughly assess the area a single molecule EhVps20t occupies at the membrane. For this, we considered the membrane of simpler compositions and took into account that the area per lipid molecule is 0.68 nm^2^ for POPC ^[32]^ and 0.55 nm^2^ for POPS ^[33]^; we assumed that the molecular areas are preserved in the mixed membrane. The ratio of labelled to unlabelled protein in the experiments (1:4) was also considered. Our approximate estimates show that the area containing a single protein molecule decreased with increasing bulk concentration of EhVps20t. For instance, the available membrane area per single protein molecule of ~102×102 nm^2^ at 125 nM of EhVps20t decreased to ~57×57 nm^2^ for the highest coverage (at 800 nM EhVps20t); see Figure 2c. For comparison, the dimensions of the protein predicted from the crystal structure of other ESCRT-III homologues are 39.2×33.7×96.4 Å^3^ (following modelling reported in ^[21c]^), but the activated protein (with open conformation) has a larger size ^[10b]^ and could prevent further protein binding by electrostatic interactions. Note that the obtained areas per single protein correspond to conditions far below surface concentrations of proteins observed to trigger curvature generation and fission due to crowding ^[34]^. Only at the highest explored concentration (1200 nM EhVps20t) were the vesicles occasionally observed to exhibit outward tubulation potentially indicating steric interactions and tendency to generate positive curvature; see inset in Figure 2b.

**Figure 2.**
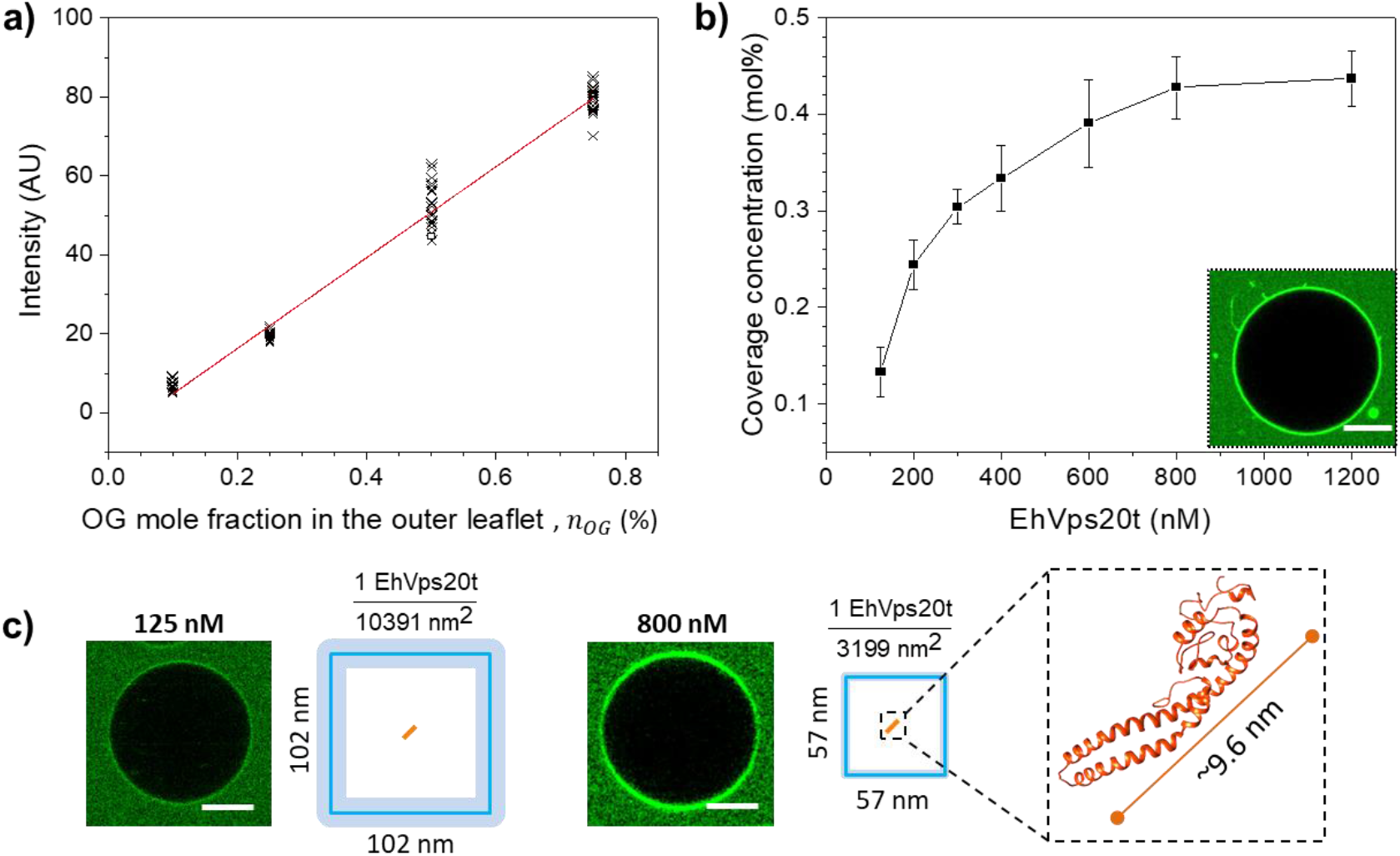
Quantitative measurement of EhVps20t bound to the membrane. (a) Calibration curve of the average intensity per pixel obtained with different concentrations of OG-DHPE in POPC:POPS (80:20) GUVs equilibrated in protein buffer (25 mM Tris, 150 mM NaCl, pH = 7.4). The symbols represent measurements on different vesicles and the red line is a linear fit (see text for expression). (b) Quantitative estimate for the coverage of EhVps20t (mole fraction with respect to the lipids) for different bulk protein concentration at which POPC:POPS:Chol:PI(3)P (62:10:25:3) GUVs were incubated. In each condition, triplicate incubations were done. Symbols represent the mean value and vertical lines the standard error of the coverage measured on 20 GUVs. The insert shows outward tubulation observed when 1200 nM of EhVps20t was used. (c) Representative images of the coverage of EhVps20t at the membrane for two different concentrations (125 nM and 800 nM), both of which are dilute and small compared to conditions of protein crowding. The depicted fractions and squares indicate the average areas occupied by a single molecule of EhVps20t (orange line, in scale with the square size; the thickness of the light blue square border illustrates the error) in the respective conditions. The predicted 3D structure and approximate size of EhVps20t in crystalline state is also given (note that upon binding the protein could unfold and occupy a larger area). Scale bars: 20 μm.

### 2.3. Stability and remodelling of intraluminal buds

Next, we aimed to explore the stability and remodelling of the intraluminal buds and how they can be affected by alterations in membrane tension and upon removal of the protein excess. Tension was modulated using two approaches: (i) osmotic inflation and (ii) micropipette aspiration. For both approaches, electroformed GUVs from the lipid mixture of POPC:POPS:Chol:PI(3)P (62:10:25:3) were incubated with 125 nM EhVps20t and 600 nM EhVps32 (mixture 1) and observed by confocal microscopy. As expected, spherical buds appeared (**Figure 3**a-c, top row of images).

**Figure 3.**
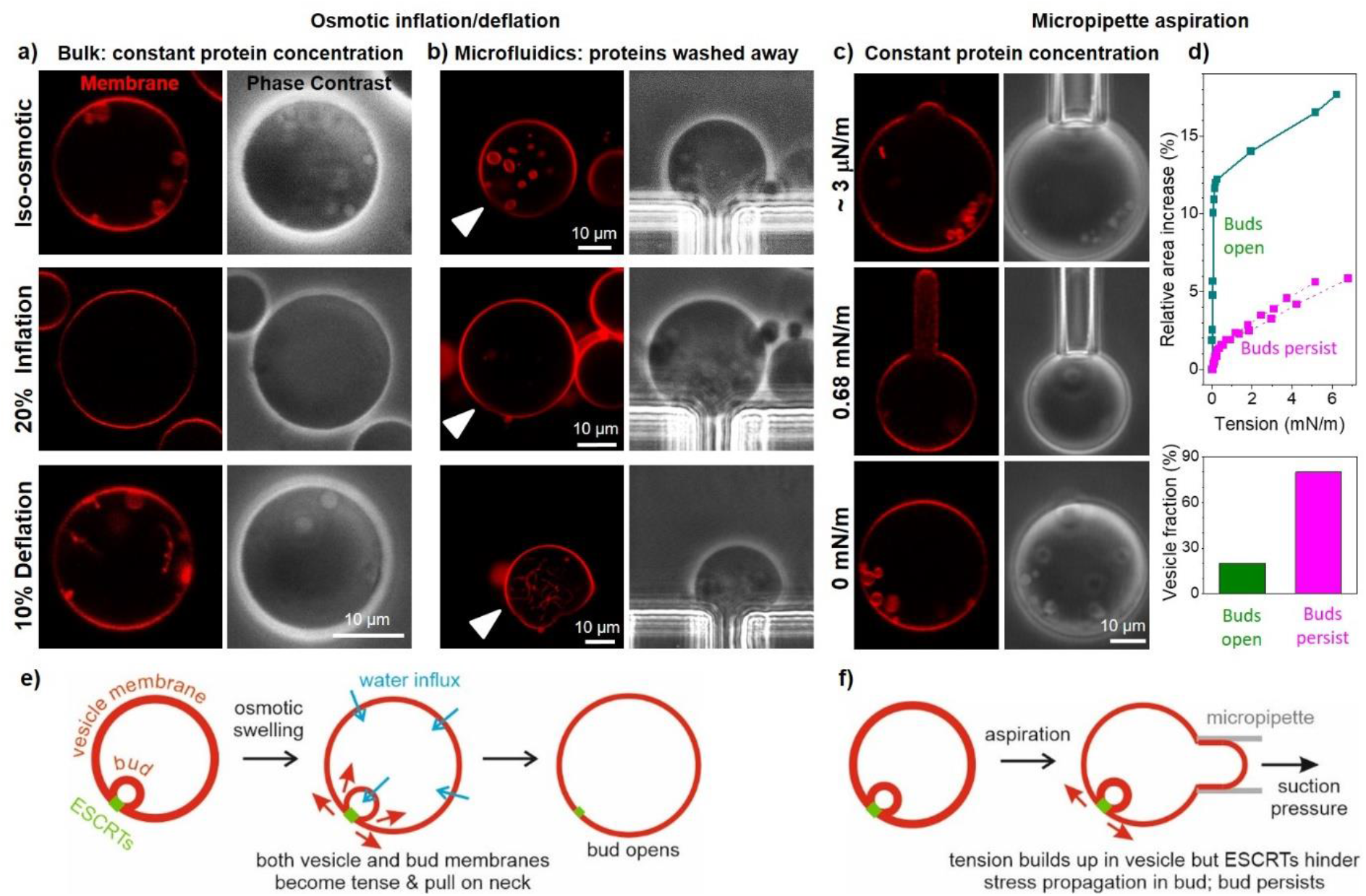
Stability and remodelling of intraluminal buds upon variations in membrane tension imposed by osmotic inflation/deflation and by micropipette aspiration of GUVs in the presence of EhVps20t and EhVps32 only. Electroformed POPC:POPS:Chol:PI(3)P (62:10:25:3) GUVs were incubated with 125 nM EhVps20t and 600 nM EhVps32 and (a) observed in bulk, (b) loaded in a microfluidic chamber or (c) aspirated in a micropipette. In all cases, intraluminal buds were observed (see first row and zoomed inset). Afterwards, (a, b) the GUVs were incubated with a hypotonic solution to inflate them up to 20% which led to bud suppression. Increase in the suction pressure in the micropipette (c, d) only occasionally (in 20% of the vesicles) led to opening of the membrane neck connecting the bud to the mother vesicle. Note that while (a) shows different GUVs, in (b) and (c) we follow the same vesicle during the experimental steps (arrowheads point to the monitored vesicle which was thoroughly examined also with 3d confocal scans). Then, a mild deflation (10%; a and b) and pressure release (c) were applied (last row of images). In the bulk experiment (a) where the total protein concentration was kept constant, we observed the formation of necklaces of small spheres (see also Movie S3), while in the microfluidic device (b), where the free proteins were washed away, tubes with sub-microscopic diameters were observed after the deflation step. In aspirated vesicles where buds open, they reform upon suction pressure release. (d) The majority of aspirated vesicles (n = 15) do not show bud opening even when the tension is increased up to about 6 mN/m above which the vesicles rupture. Three examples for area-tension traces are shown. (e) During osmotic inflation, water permeates across the membranes of both the mother vesicle and the bud, increasing the tension in both membranes (two pairs of red arrows); both compartments are drawn with thinner contours to reflect the higher tension of their membranes; indeed, in the high tension regime of stretching, the membrane should become thinner). This leads to opening of the bud neck. (f) On the other hand, by micropipette aspiration only the membrane tension in the mother vesicle (only one pair of red arrows) is directly increased; only the vesicle membrane becomes tense as reflected by the thinner contour, but not that of the bud. Stress propagation to the bud membrane is plausibly hindered by the ESCRT proteins assembly stabilizing the bud neck.

#### Osmotic inflation

For this approach, the mixture of proteins and GUVs (mixture 1) was diluted 1:2 with a hypotonic solution (of 20% lower osmolarity) to inflate the vesicles, see SI Section S1. After inflation, we observed that the previously formed intraluminal buds disappeared, implying that the bud necks could be opened by increasing the membrane tension (Figure 3a, inflation). We then performed a mild deflation step of 10% osmolarity increase. In this case, we detected the formation of buds and long necklaces of small spheres with a typical size of 1-2 μm, Figure 3a, deflation (see also Movie S3). Note that a constant concentration of proteins was maintained throughout the whole experiment; in the absence of proteins or when the total amount was diluted, such necklace-like structures were not observed upon deflation.

In these bulk experiments, the initial state of the observed vesicles (and in particular, whether the vesicle had internal/external structures before the inflation/deflation steps) is not known. To overcome this uncertainty, microfluidic devices were used to follow the same vesicle exposed to inflation and deflation. In this approach, the unbound proteins are removed and some bound proteins may also be washed away (note that experiments with constant concentration of proteins was not feasible in this case because of the high protein amounts required). GUVs (mixture 1) were loaded in a microfluidic chamber. The trapped vesicles typically exhibit a smaller number of buds than those in bulk measurements because the posts of the traps provide additional constraints on the trapped vesicles reducing the excess area available for budding (note that upon stopping the flow, the deformation from the posts is absent but budding is preserved). After a 20% inflation step, the GUV volume increased and the buds disappeared (Figure 3b, inflation), similarly to the behaviour of the vesicles under bulk dilution (Figure 3a, inflation). However, upon 10% deflation, the vesicles did not develop micron-sized buds but only inward tubes with sub-optical diameters (Figure 3b, deflation) presumably resulting from the solution asymmetry. This suggests that in the bulk experiment (Figure 3a), where the proteins are still present (contrary to the case of deflation in the microfluidics experiment as in Figure 3b), the newly formed buds and necklaces, whose micron-sized buds are comparable in diameter to those of the intraluminal buds, are stabilized by newly-bound proteins available in the external solution (but absent in the microfluidics experiment). The stability of single buds versus interconnected necklace multi-buds depends on membrane spontaneous curvature^[35]^, here governed by the proteins; both single buds and necklaces can be formed for the same shape parameters, i.e., for the same spontaneous curvature and vesicle volume-to-area ratio, see ^[35c]^, but the kinetics and dynamics of these parameters likely determine the stable shape. Note that studies in worms^[36]^ and in plants^[37]^ showed similar buds interconnected into necklaces (concatenated structures) forming under the influence of ESCRTs.

In both, bulk and microfluidics experiments, the intraluminal buds are suppressed by osmotic inflation (they open up as a result of built-up membrane tension) and do not reform to the same extent upon tension release (deflation) suggesting irreversible remodelling of the scaffold-like structure of polymerized protein. Even though the binding of EhVps32 to EhVps20t is relatively strong ^[38]^ depleting EhVps32 from the bulk could result in desorption of the protein. We thus speculate that in the microfluidic chamber, upon increased tension and partial protein desorption, the EhVps32-protein scaffold irreversibly deforms failing to trigger the bud-like invagination when the membrane is deflated again. In contrast, in the bulk experiments, the free proteins in the bulk solution can bind triggering the formation of new inward structures with dimensions similar to the initially observed intraluminal buds.

#### Micropipette aspiration

As we mentioned above, adding proteins together in a mixture or in a step-by-step manner does not have any influence on the number of GUVs with ILVs (Figure S7). To isolate the effect of potentially desorbing proteins during the microfluidic inflation experiments, we probed the bud stability while keeping the protein concentration constant. For this we used micropipette aspiration, see Section S3 in the SI. A small fraction of the vesicles showed bud opening (Figure 3c,d). One such example is shown in Figure 3c. The tension was gradually increased and the buds opened at tensions slightly above 0.01 mN/m. After gradual decrease in the tension, the buds reformed. The majority of the aspirated GUVs (80%) did not exhibit bud opening even in the regime of high membrane tension up to around 6-7 mN/m, close to the lysis tension. Above this tension, the vesicles collapsed being sucked up inside the pipettes. When bud opening was observed during aspiration (20% of the vesicles), it occurred at tensions (0.01-0.1 mN/m) which are considerably higher than the spontaneous tension (generated by the spontaneous curvature arising from the protein adsorption), the upper limit of which is on the order of 4×10^-5^ mN/m as estimated from the intraluminal bud size, see Section S3 and Refs. ^[35a, 39]^. Thus, protein assembly and scaffolding must be stabilizing the buds. For a radius *R_ne_* of the membrane neck in the range between 5 and 25 nm, a tension of Σ = 0.1 mN/m at which neck opening was observed, implies the neck opening force *f*~2π*R_ne_*Σ~ 3 ÷ 16 pN generated by the membrane tension. This argument however ignores the membrane tension in the bud membrane and assumes uniform curvature-elastic parameters, the validity of which is unclear.

### 2.4 Protein domains, and protein and lipid mobility in intraluminal buds

To check whether the assembly process of the two ESCRT proteins can be visualized in real time in the membrane of the GUVs, lower concentrations of the protein were used. For the protein concentrations discussed so far, we observe that budding has occurred in most vesicles already after less than 5 min (Figure S8a). At lower EhVps32 protein concentrations, bud formation is slowed down. For instance, at 300 nM of EhVps32, maintaining EhVps20t concentration at 125 nM, the budding is delayed by up to 15 minutes after protein addition. Furthermore, when we increased the concentration of EhVps32 to 750 nM, the budding process started already 1 min after protein addition. In accordance with these observations, the number of buds gradually increases with the concentration of EhVps32 (see Figure S8). After 10 min incubation of GUVs with 125 nM EhVps20t and 300 nM EhVps32, we were able to detect EhVps32-rich protein domains in the membrane by monitoring the fluorescently labelled analogue OG-EhVps32 (top row in **Figure 4**a, arrow). It is important to note that in the presence of EhVps24 we did not detect protein domain formation (probably because of the high protein concentration or the enhancing activity of EhVsp24 that changes EhVps32 filaments conformation, while speeding up the process), suggesting a regulatory role of this protein. Interestingly, EhVps32-rich domains appeared to differ in their lipid composition as well, as concluded from the enhanced intensity of the membrane dye DiIC_18_ (top row in Figure 4a). We cannot exclude that these domains represent accumulation of membrane folds with sub-optical resolution dimensions, however, no thickening of the vesicle membrane was detected under phase contrast (top and middle row in Figure 4a). Fifteen to twenty minutes after the incubation (throughout which the vesicle and the domain were constantly monitored), membrane invagination and budding occurred at the site where the protein domain was formed (middle row in Figure 4a, arrowhead, Figure S9). The formation of the intraluminal bud was associated with partial to complete dissolution of the EhVps32-rich domain (bottom row in Figure 4a). Presumably, the accumulated protein rearranged while scaffolding the inward bud. Note that the signal from the domain-segregated lipids (red channel in bottom row of Figure 4a) did not fully decay even after the formation of the intraluminal bud. However, EhVps32 domains were detected also in unlabelled GUVs (Figure S9), suggesting that their formation is mediated mainly by this protein.

**Figure 4.**
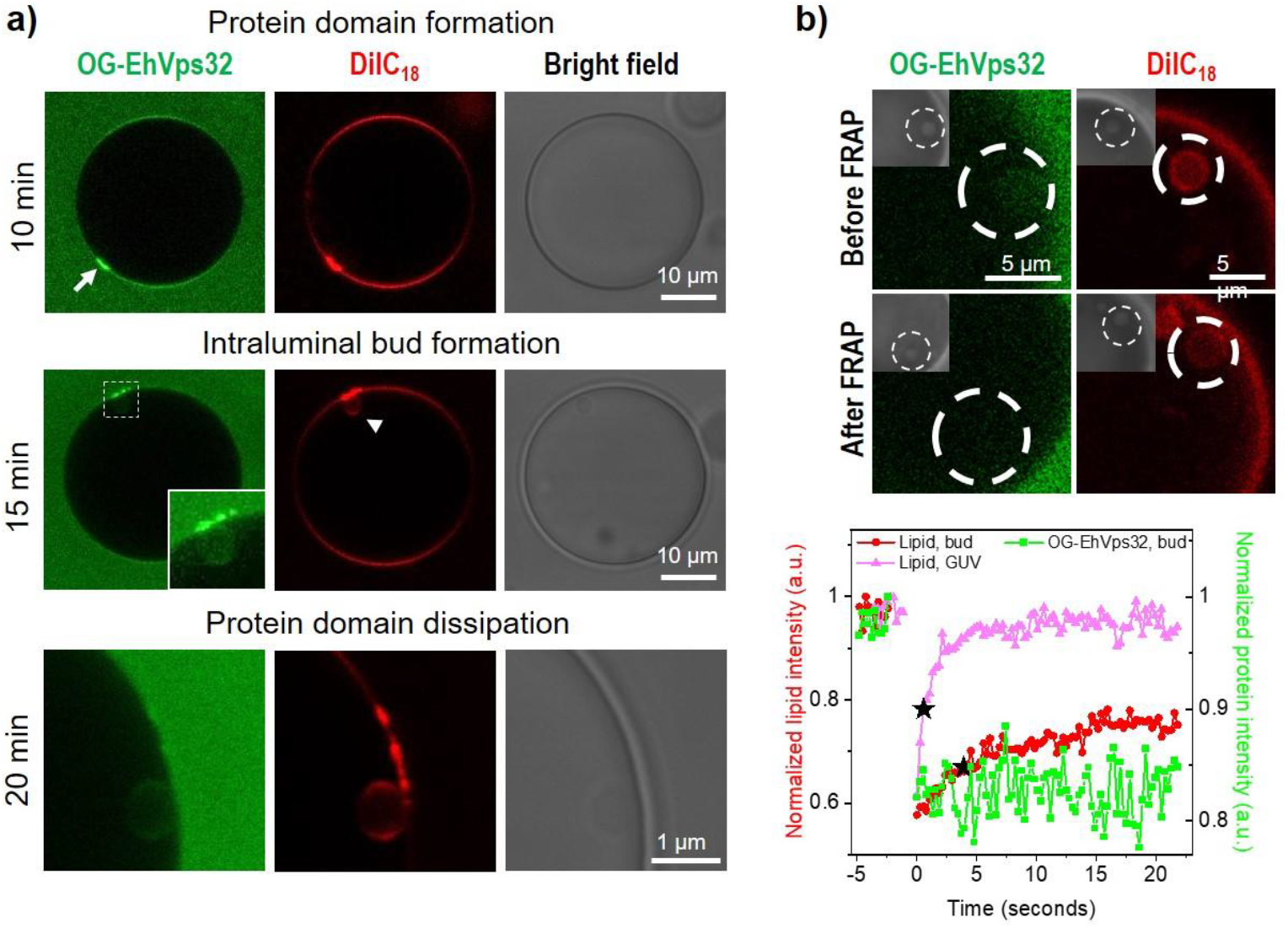
EhVps32–enriched domain formation and protein and lipid mobility in intraluminal buds. GUVs composed of POPC:POPS:Chol:PI(3)P (62:10:25:3) were incubated with 125 nM of EhVps20t and 300 nM of EhVps32 (20% of EhVps32 was labelled; both proteins are added at the same time). (a) Images of the same GUV taken after incubation of 10, 15 and 20 minutes (inset and last row showing the zoomed region of the same intraluminal bud) respectively showing the formed protein-enriched domain, the formation of intraluminal bud and protein domain disassembly. (b) FRAP measurements made on an intraluminal bud formed on a different vesicle, of which only a zoomed segment is shown with the bleached region indicated by the dashed circles. The images show protein and lipid fluorescence from the region of the intraluminal bud before (top row) and after (bottom) bleaching; inserts display the phase-contrast image clearly showing the location of the bleached intraluminal bud. The curves below (collected consecutively) demonstrate the lack of fluorescence recovery of the protein signal (green) and partial recovery of the lipids in the bleached intraluminal bud (red); the pink curve represents lipid recovery signal in a region at the vesicle surface. The black stars indicate the half time of the lipid fluorescence recovery, which is 0.57s and 3.87s for the vesicle surface and the intraluminal bud, respectively.

To resolve whether the protein remained locally immobilized at the membrane of the intraluminal bud indicating a rigid assembly, we examined the recovery of fluorescence signal from OG-EhVps32 after photobleaching, see part on fluorescence recovery after photobleaching (FRAP) in SI Section S1. The FRAP experiments (Figure 4b) did not indicate recovery of the OG-EhVps32 signal after bud bleaching suggesting that the assembled proteins are immobilized. Note however that the initial signal in the bud is generated both from bound and unbound protein, which cannot be distinguished. We thus conclude that EhVps32 is locally immobilized and is not able to pass through the narrow neck of the bud nor is additional protein from the mother vesicle able to diffuse through the neck (only a limited amount of lipid does). In contrast, after photobleaching the same intraluminal bud, the lipid fluorescence partially recovered (contrary to observations for the human Vps2 protein, CHMP2B, which was reported to prevent lipid diffusion ^[40]^). It is worth noting that full recovery was observed when photobleaching a region of the lipid membrane in the top area of the GUV (Figure 4b), with a lipid diffusion coefficient 5.8 μm/s which is comparable to literature data. Presumably, the EhVps32 scaffold impedes the lipid mobility within the membrane of the bud.

### 2.5. Membrane remodelling by EhVps20t and EhVps32 is antagonistic

Next, we aimed at understanding the mechanisms controlling the size of the generated buds. Precise imaging and size determination of buds is hindered by the vicinity of the mother GUV membrane. We thus assessed the sizes of ILVs. They were measured at 125, 300 and 600 nM concentrations of EhVps20t. Each EhVps20t condition was tested in combination with three different EhVps32 concentrations, arbitrarily named low, medium and high (300, 600 and 1000 nM respectively), all in the presence of 200 nM EhVps24 necessary for membrane scission. All proteins were added simultaneously to the vesicle suspension. Solution osmolarities were carefully adjusted (see SI Section S1); note that incubation with protein-free buffers led only to the vesicle deflation but no formation in micron-sized buds/ILVs. Based on previous work ^[21b, 21c]^, EhVps32 concentrations lower than 300 nM were not sufficient to generate ILVs in giant vesicles, and concentrations higher than 1.3 μM induced GUV disruption (presumably resulting from high steric surface pressure as observed with other proteins ^[34]^). All possible combinations of both protein concentrations were tested and the size of the ILVs was measured from 3D confocal scans. The diameter of the ILVs was found to increase with EhVps20t concentration in the batch (**Figure 5**a), presumably because of membrane stiffening. On the contrary, EhVps32 appeared to stipulate higher curvature resulting in smaller size of the ILVs with increasing protein concentrations (Figure 5a). Taken together, these results suggest that the two proteins act antagonistically and that at least two different mechanisms influence the curvature of the membrane and therefore, the size of the generated ILVs.

**Figure 5.**
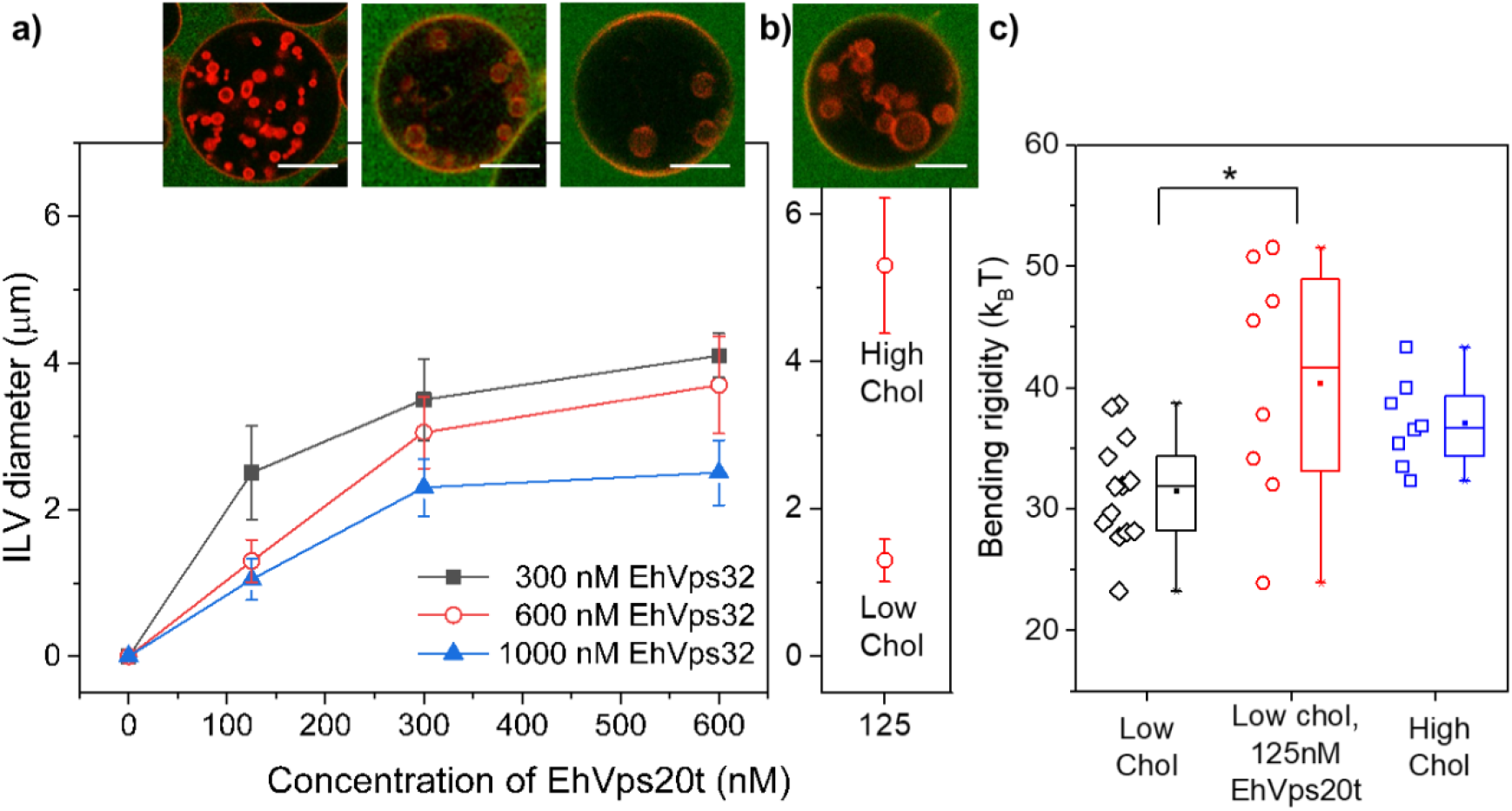
Antagonistic effects of EhVps20t and EhVps32 on the size of ILVs and role of cholesterol content in the membrane. (a) The ILV size was measured on GUVs made of POPC:POPS:Chol:PI(3)P (62:10:25:3) in protein buffer (25 mM Tris, 150 mM NaCl, pH = 7.4). Three different concentrations of EhVps20t (125, 300 and 600 nM) were tested in combination with three concentrations of EhVps32 (300 nM in black, 600 nM in red and 1000 nM in blue). All proteins were added at the same time. (b) For high-cholesterol fractions in the membrane, GUVs made of POPC:POPS:Chol:PI(3)P (52:10:35:3) were used. In all conditions (a, b), the concentration of EhVps24 was maintained at 200 nM. Proteins were added simultaneously to the vesicle suspension. The diameter of ILVs generated in all possible combinations, was measured and plotted against the concentration of proteins. Experiments were performed by triplicate and in each condition 20 GUVs with at least four ILVs were measured. The data represent the mean and the standard error. The confocal sections on top display representative images of the red open-circle data in panel (a) and high-cholesterol composition in panel (b); in here, 20% of labelled EhVps20t (green) was used to follow the effect on the membrane (red). Scale bars: 10 μm. (c) The bending rigidity of the membranes with low and high cholesterol fractions, POPC:POPS:Chol:PI(3)P (62:10:25:3) and (52:10:35:3) respectively, was measured from fluctuation spectroscopy of GUVs prepared in 20 mM sucrose and diluted in isotonic glucose solution with or without EhVps20t (125 nM). The measurements (see Materials and Methods) were conducted at room temperature (~ 23°C). The data points on the left of each bar show the individual measurements on different vesicles (at least 8 vesicles per composition were examined). Mean values and standard deviations are also given.

### 2.6. The size of ILVs is controlled by competing mechanisms of curvature generation and regulation

In the following, we attempted to disentangle the contribution of the ESCRT-III proteins in regulating the size of the ILVs. We considered only the effect of EhVps20t and EhVps32 as EhVps24 appears to induce the scission of vesicles from formed intraluminal buds with already defined diameter. Previous work demonstrated that EhVps32 and its activated version, EhVps32(1-165) do not bind to the membrane of negatively charged GUVs and do not produce ILVs in the absence of EhVps20t ^[21c]^. Therefore, EhVps32 by itself is unable to generate significant changes in the bare membrane of GUVs with endosomal composition when compared to incubation with buffer only. As observed in the above experiments, increasing the concentration of EhVps20t added to GUVs, increased the size of the generated ILVs, while the opposite effect was observed with EhVps32, see Figure 5a, with trends suggesting that EhVps32 builds its effect on the EhVps20t-coated membrane.

To distinguish the effects of the two proteins and their bending energy contributions, we first probed whether the bending rigidity of the GUVs is altered by the adsorbed EhVps20t. For this, we performed fluctuation spectroscopy^[41]^ of vesicles in sugar solutions (see Materials and Methods); the coverage of EhVps20t on the membrane in these conditions was found to be the same as that in the salt buffer (Figure S6). Note that measurements in the presence of EhVps32 were not feasible because the formed inward buds suppressed the fluctuations. The bending rigidity of protein-free GUVs (31.5±4.5 k_B_T, the value is relatively high compared to that of fluid neutral membranes but consistent with increased stiffness observed for membranes with higher charge density^[42]^) was found to increase (to 40.4±10.0 k_B_T) when 125 nM of EhVps20t was introduced (Figure 5c). Here, k_B_T is the thermal energy at room temperature. The observed stiffening cannot be caused by protein myristoylation because EhVps20 lacks the glycine residue in its N-terminal (see Figure S10), which would be necessary for the myristolylation process. Higher protein concentrations (200 nM and higher) could not be explored because the shape fluctuations of the membrane were suppressed and could no longer be analysed; we also observed “protein clusters” at the membrane of the GUVs (Figure S11), which are absent at high salinity. These clusters affected the detection of the vesicle contour for bending rigidity measurements. The membrane stiffness was found not to be influenced by the salinity buffer itself, Figure S12.

To find out whether the rigidity of the protein-free membrane can regulate the diameter of ILVs in the same way as EhVps20t does, we measured their size in GUVs with stiffer membranes of increased cholesterol fraction. Note that cholesterol is known to increase the bending rigidity of some ^[43]^ but not all membranes ^[41a, 44]^, see overview in ^[45]^. To confirm that we work with stiffer membranes, we measured the bending rigidity of the vesicles with low and high cholesterol fractions. As a high-cholesterol membrane we explored vesicles made of POPC:POPS:Chol:PI(3)P (52:10:35:3), which compared to the standard mixture (62:10:25:3) preserves the surface charge (to ensure similar binding of the proteins) while increasing the cholesterol fraction by 10 mol%. The respective membrane bending rigidities for the high-cholesterol mixture and the standard mixture with low cholesterol content were found to be 37.0±3.6 k_B_T and 31.5±4.5 k_B_T (Figure 5c). Consistently, for the stiffer membrane, the ILV size was significantly larger, see Figure 5b, suggesting that it is modulated by changes in the membrane bending rigidity imposed by the adsorption of EhVps20t. Protein density variation affecting the ILV size in vesicles with different cholesterol fraction can be excluded as the protein coverage for both membranes was found similar (Figure S6). Note also that for much stiffer membranes made of DOPG:eSM:Chol 20:60:20 no intraluminal bud formation was detected, confirming that the stiffness of the bare membrane plays a significant role.

Changes in the membrane stiffness are correlated to changes in the spontaneous curvature which is reciprocal to the bending rigidity ^[46]^. In homogeneous membranes, the magnitude of the spontaneous curvature is roughly inversely proportional to the bud size ^[47]^, see also section 4 in Ref. ^[48]^ for a summary of approaches to assess the membrane spontaneous curvature. In our experiments, a decreased ILV size was observed when increasing the EhVps32 concentration (at a fixed EhVps20t concentration), suggesting the generation of more negative spontaneous curvature or the formation of a polymerized protein scaffold at the membrane. The overall impact of the two proteins on the membrane with endosomal composition is summarized in **Figure 6a** and discussed further below. The final ILV size is governed by a competition between the bending rigidity, increased by EhVps20t binding, and the spontaneous curvature enhanced by EhVps32 polymerization.

**Figure 6.**
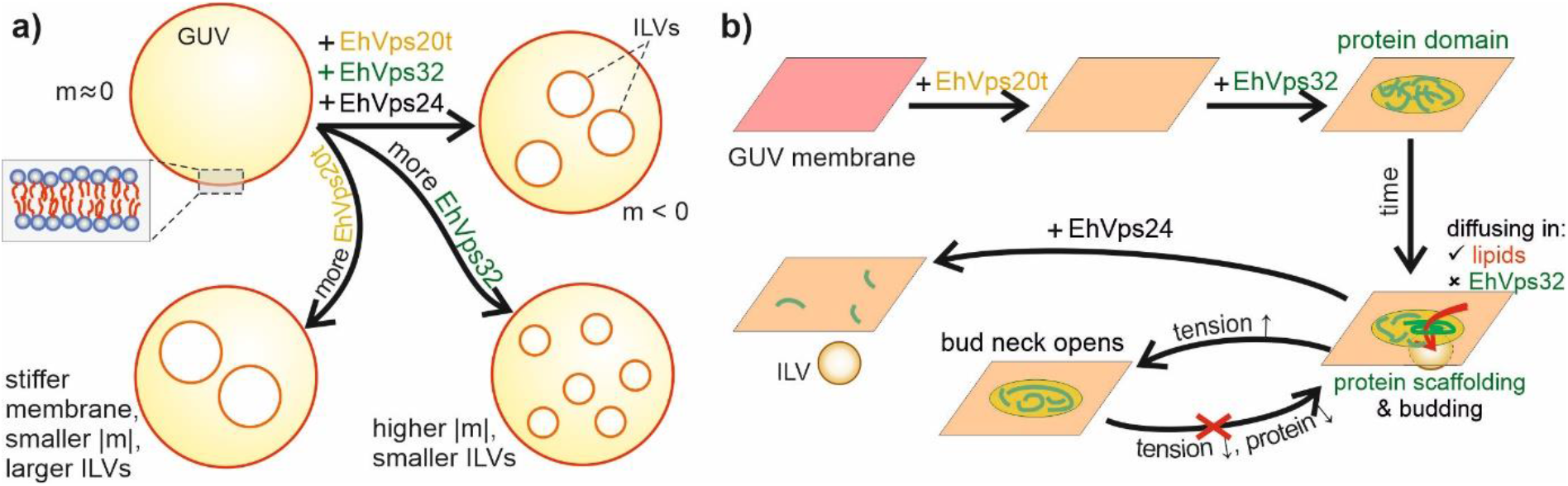
Schematic illustration and proposed scheme of the action of ESCRT-III proteins on membranes of endosomal mimetic. (a) EhVps20t and EhVps32 influence the spontaneous curvature, m, and bending rigidity of the GUV membrane, regulating the size of intraluminal vesicles. Increasing the concentration of EhVps20t leads to membrane stiffening and larger ILV size while raising the concentration of EhVps32 can enhance the membrane curvature while scaffolding the membrane into smaller ILVs. (b) Proposed schematics of action. EhVps20t homogeneously binds to the membrane, while EhVps32 generates protein-rich domains. Over time, a nascent vesicle (intraluminal bud) is formed at the site of the domain. Protein diffusion through the neck is presumably blocked but lipids can still diffuse in the bud although to a limited extent. The bud neck can be open upon tension increase, but releasing this tension does not allow for reforming this bud supposedly because of the irreversibly distorted protein assembly. Abscission of the intraluminal bud is mediated by EhVps24.

## 3. Discussion

In general, fission implies a sequence of membrane deformation processes that finally lead to the formation of two independent membrane compartments from a single one. In some cases, fission is mediated by membrane protein coats or complexes and constriction (i.e., dynamin). In others, it is triggered by local membrane perturbation (i.e., by lysolipids and ENTH) ^[49]^. Recently, another mechanism has been discovered where molecular crowding seems to be sufficient to drive membrane scission as long as proteins bind to the membrane at a high coverage and their steric pressure overcomes the barrier for membrane scission ^[34]^. Even at low protein concentrations, spontaneous curvature can generate forces sufficient to constrict and fission the membrane ^[35a, 50]^. Until now and despite the effort, the precise mechanism by which the ESCRT machinery triggers membrane fission is not completely understood. Over the past few years, several studies have reconstituted the action of the ESCRT-III machinery in GUVs ^[16a, 20e, 21, 23, 51]^. In most of these experiments, purified ESCRT-III proteins were added to a batch of GUVs contained in a chamber and images were taken of different GUVs. The proteins remained in the batch throughout the whole reaction making it difficult to assign a specific task to each protein. In this study, using microfluidic technology we were able to trap GUVs and follow the ESCRT-induced remodelling on the same vesicle. The main advantage of this approach is that unbound proteins are washed away prior to the addition of the next protein allowing a more controlled experiment and decoupling the distinct roles of the different proteins. In this way, we show that using a minimal system of only three ESCRT proteins and a microfluidic approach, we are able to establish a synthetic microcompartmentalized cell mimetic, in which the compartment size and its content can be controlled. The three-protein construct also offers a relatively simple approach for establishing synthetic cell division ^[52]^, even in the absence of an energy source such as ATP. Three main events during the ESCRT-III-mediated membrane remodelling have been identified using a single-vesicle assay and elucidated: binding of EhVps20t to the GUV membrane, invagination and budding of this membrane triggered by EhVps32 and membrane scission after EhVps24 addition (Figure 1), thereby mimicking the *in vivo* production of intraluminal vesicles with a minimal set of components. It is important to mention that the micrometer size of ILVs in our reconstitution experiments is significantly larger than that of ILVs observed in cells (~50 nm^[53]^). One possible reason as proposed previously^[54]^ is the higher specific binding efficiency of ESCRT-III to the endosomal membrane compared to the GUV membrane as well as the absence of cargo proteins in the latter system. However, we did observe that the generated buds were homogeneous in size, which suggest that the size regulation is governed by factors such as protein surface coverage and membrane bending rigidity as demonstrated here (Fig. 5).

It has been suggested that ESCRT-III is a dynamic polymer where multiple subunit turnover events trigger the deformation and scission of the membranes. In particular, Pfitzner et al. ^[55]^ demonstrated that the Vps32 polymer recruits the Vps2-Vps24 sub-complex, which in turn recruits Vps4. The enzyme promotes polymer growth and exchanges Vps24 for Did2, stimulating Vps4 activity, which leads to the disassembly of Vps32 and Vps24 filaments. Finally, the increasing levels of Did2 promote 3D deformation and intraluminal budding, after which Ist1 induces final scission ^[55]^. In our model of GUVs system, we have found that EhVps4 was not necessary for the formation of ILVs. Protozoan parasites, including *E. histolytica* exhibit a substantial reduction of the ESCRT machinery compared to higher eukaryotes such as yeast and human. Indeed, ESCRT-III subunits in this parasite lack MIT-interacting motifs (MIM) which are responsible for the binding to microtubule-interacting and transport (MIT) domain of Vps4 in its human and yeast orthologue ^[56]^. Therefore, we hypothesize that the mechanism of subunit turnover responsible for membrane remodelling and budding as well as ESCRT-III mediated scission *in vivo* could be occurring (at least in part) independently from EhVps4 as we show in our work.

To increase the understanding of the mechanisms that drive the membrane remodelling processes, we focused on investigating the first two steps (binding and invagination), which also define the size of intraluminal vesicles. We first measured the coverage of EhVps20t on the membrane of the GUVs (Figure 2) and observed saturation of the protein surface concentration at the membrane when the bulk protein concentration was about 800 nM. Presumably, EhVps20t acts as a nucleation point for the polymerization of EhVps32 as previously suggested ^[57]^, therefore providing the specific surface area that would control the size of the generated ILV. Our estimates for the area occupied by EhVps20t (Figure 2c) agree with the diameters observed in deep-etch electron microscopy of plasma membranes from cultured cells depleted of Vps4 where ESCRT filamentous assemblies with diameters between 108 ± 30 nm were preserved ^[58]^. We conclude that EhVps20t restricts the area available for EhVps32 polymerization and, therefore, ILV size regulation. From a synthetic point of view, ILVs represent microcompartments that are biomimetic analogues of cellular organelles and having control over their size and composition is advantageous. Obviously, the membrane coverage and bulk concentration of these two proteins could provide control parameters for organelle size in artificial cells, at least in the range 1 – 6 μm as shown in Figure 5a,b. Multicomponent membranes prone to phase separation could ensure a distinct composition and phase state of the ILV membrane (different from that of the GUV)^[31]^. Furthermore, the microfluidic approach introduced here allows the subsequent loading of these compartments with solutions different from that in the vesicle interior/exterior, thus offering a pathway for performing localized and sub-compartmentalized processes such as protein synthesis or enzymatic reactions in a cell-like environment.

How easy it is to bend a flat membrane, for example when forming invaginations or outward buds, depends on the bending rigidity. The bending energy required to form a spherical bud from a flat membrane (of negligible spontaneous curvature) is 8*πκ*. In the absence of an external pulling force, the direction of budding is defined by the membrane spontaneous curvature ^[46a]^. In the presence of EhVps20t, the vesicle membrane becomes stiffer (Figure 5c) but it does not exhibit buds and the spontaneous curvature remains close to zero. The latter curvature should then be induced by EhVps32. A number of mechanisms exist for generating and regulating the spontaneous curvature of membranes ^[46a, 48, 59]^. Both the bending rigidity (in heterogeneous membranes) and the spontaneous curvature can control the membrane shape (even to the extent of triggering membrane scission in membranes with fluid domains ^[50, 60]^). But these two membrane properties are often interrelated: the spontaneous curvature induced by adsorption or depletion layers, for instance, is inversely proportional to the bending rigidity ^[46a]^ and thus, stiffer membranes would result in larger ILVs (i.e. would exhibit a spontaneous curvature which is lower in absolute magnitude). Entropic and steric effects between externally adsorbed proteins are typically expected to increase the magnitude of the spontaneous curvature of positive sign (i.e. the membrane tends to bulge more strongly towards the compartment containing the protein) with increasing coverage. Thus, adsorption of EhVps20t should intuitively lead to decreasing the magnitude of negative spontaneous curvature, which is consistent with the increase in ILV size (Figure 5). Thus, EhVps20t decreases the curvature magnitude (i) by entropic interactions and/or (ii) by effectively increasing the membrane bending rigidity.

Our results show that the size of the intraluminal buds is controlled not only by the EhVps20 concentration but also by the amount of EhVps32 present in the bulk whereby the two proteins influence the ILV size antagonistically (Figure 5). To be able to resolve the process of ILV formation, we slowed down the reaction by reducing the concentration of EhVps32 (to 300 nM). This approach allowed us to observe the formation of EhVps32-rich domains and the subsequent formation of membrane buds in the region of these domains (Figure 4a), which has not been reported previously. Scission mediated by EhVps24 could result from reorganization of EhVps24 into assemblies that narrow the neck of the bud, as demonstrated for Vps24-induced Vps32 (Snf7) helical assemblies in Ref. ^[16a]^. Although ATPase activity by Vps4 may be essential for regulating the dynamic behaviour of Vps32 filaments, here, we observe that EhVps32 has an intrinsic ability to self-associate forming homopolymers^[61]^ (without the requirement of additional factors such as Vps4) as observed for the Vps32 homologue in *Caenorhabditis elegans* ^[62]^. The minimal system of three ESCRT proteins as used here generates intraluminal vesicles in the absence of ATP, which provides a new and simpler pathway to vesicle division without the need to be concerned about any chemical energy supply. As recently demonstrated^[54]^, ESCRT-mediated budding in human cells is also a passive process driven by crowding of upstream ESCRTs (−0, -I and –II) coupled to a steep decrease in Gaussian curvature. The accumulation of upstream ESCRTs leads to recruitment of ESCRT-III/Vps4, which in turn triggers neck constriction and scission. Furthermore, in cells depleted of ESCRT-III, even though the overall ILV generation was impaired, their formation was still observed albeit at a lower degree, promoting the hypothesis that upstream ESCRTs initiate the membrane budding process through protein crowding^[54]^ (similarly to previously reported effect of curvature modulation by GFP binding to membranes^[63]^) even though, after forming a bud neck, scission can be achieved also at low protein coverage^[50]^. It would be interesting to investigate the structure of the EhVps32-rich domains with high-resolution techniques. These micron-sized domains appear to be transient and disassemble after the buds were fully formed (Figure 4a, bottom row), suggesting lateral mobility of the proteins bound to the vesicle surface. However, no diffusion of the membrane-bound proteins between the mother membrane and the buds could be detected in the FRAP measurements on bleached intraluminal buds (Figure 4b). In contrast, the underlying lipids partially recovered (Figure 4b). We speculate that the protein assembly around the membrane neck between the bud and the mother membrane as well as the anchor points of the protein assembly in the bud act as obstacles slowing down and obstructing lipid diffusion.

As shown in Figure 3, the buds open during osmotic inflation, while they do so only in a small fraction of vesicles when membrane tension is applied by micropipettes. The osmotic pressure acts to inflate and stretch the outer vesicle membrane and, as water permeates first through the vesicle membrane and then through the bud membrane to balance the osmotic differences. As a result, the bud gets inflated and its membrane expands/stretches as well ultimately leading to neck opening (note that the bud neck is relatively small to allow fast influx of solution to balance the osmolarity difference and the latter is mainly balanced through transmembrane permeation). Thus, the stress applied on the bud neck via osmotic inflation is imposed both from the increasing tension of the mother-vesicle as well as that of the bud membrane both pulling on the neck, see Figure 3e. The osmotic swelling of a 20 μm vesicle is established already in less than a second if we consider membrane permeability of about 15 μm/s ^[64]^, and thus the tension in the membrane of intraluminal buds builds up almost immediately. In the case of aspirated vesicles, on the other hand, the tension is directly applied only to the mother-vesicle membrane while stress propagation to the bud might be impeded by the protein assembly in the bud neck region, see Figure 3f, which is consistent with the lack of recovery in the FRAP measurements. Further increase in the mother-vesicle tension then typically leads to vesicle rupture before the bud can open. This explains the difference in the observations that under osmotic inflation buds open, while they are unlikely to do so with a rapid increase of the suction pressure in the pipette. These findings indicate that experiments on bud stability as a function of membrane tension typically performed via osmotic swelling in the bulk (which is also easier than micropipette aspiration) might point to misleading conclusions regarding the effect of tension. Osmotic swelling experiments are also presumably less relevant considering that cells are rarely exposed to a large osmotic shock. Considering the obtained results, the process of ILV formation in membranes mimicking endosomal composition as explored here is summarized in the sketch proposed in **Figure 6b**: EhVps20t homogeneously binds to the membrane (as shown in Figures 1, 2), while EhVps32 generates clusters appearing as protein-rich domains (a few microns in size) at the vesicle surface. Over time, at the location of the domain, a nascent vesicle (bud) is formed (Figure 4a), which detaches only in the presence of EhVps24. Increase in the membrane tension over the vesicle and the bud (as done with osmotic inflation) opens the bud distorting the scaffold formed by EhVps32. Upon tension release, new buds may reform only if previously unbound protein is present in the bulk (Figure 3). For slightly stiffer bare membranes, the mechanism is similar, however leading to larger bud and ILV size as in Figure 5. Our results demonstrate the competing roles of bending rigidity, which at low values facilitates closing the bud neck, and membrane tension, which acts in the opposite direction, namely to open the neck.

## 4. Conclusion

In conclusion, we propose that the mechanism of ESCRT-III mediated scission starts with the binding of EhVps20 proteins to the membrane which act as nucleation sites for EhVps32 recruitment. In addition, EhVps20 increases the membrane stiffness which competes with an increment in the spontaneous curvature triggered by EhVps32 incorporation. The balance of both forces produces intraluminal buds of various sizes depending on the concentration of the two proteins. The buds are stabilized by EhVps32 scaffolds (with immobilized protein but partially mobile lipids), which can be distorted upon increased membrane tension leading to bud opening.

Before an ESCRT-generated bud is released from the mother vesicle, it must be connected to this vesicle via a closed membrane neck. In general, such a closed neck can be stabilized by spontaneous curvature, scaffold adhesion, or line tension of an intramembrane domain boundary ^[35a, 39]^. So far, we have not been able to draw reliable conclusions about the relative importance of these different stabilization mechanisms, which remains an important challenge for future studies.

We demonstrated that a minimal set of only three ESCRT-III proteins is sufficient for fission of membrane necks. This finding is crucial for minimalistic approaches in synthetic biology aiming at reconstitution of cell division (with a minimal divisome) ^[50, 52]^. We also showed that the size of ILVs is governed by the protein concentration and membrane bending rigidity. These two factors offer a route for controlling the size of intracellular organelles in artificial cells. The number of organelles (ILV) would depend on the area-to-volume ratio of the initial cell (GUV). While microfluidic techniques for the production of nested vesicles in vesicles (vesosomes) allow direct mechanical control over the size of the different compartments (e.g. via adjusting flow pressure and chip geometry), membranes created from double emulsions and/or oil/water phase transfer are not suitable for the reconstitution of proteins such as ATPases and other membrane enzymes because of the inherent leaflet-by-leaflet assembly of the membrane and the presence of oil. In contrast, the strategy of controlling compartment size via the interplay of ESCRT proteins and composition (modulating the membrane bending rigidity) is closer to nature and might offer new routes towards generation of smart synthetic cells. With the microfluidic technology used here, one is also able to load the different compartments with different solutions in a stepwise manner thus allowing for localized and compartmentalized processes as in cells. Presumably, establishing liquid ordered – liquid disordered phase separation in the membrane will allow to control also the compartment membrane composition with ILVs budding preferably from one type of membrane domain ^[31, 60]^. In the field of synthetic biology, microcompartmentalized vesicles are key to reverse-engineering of eukaryotic cells with reconstituted functionality.

## 5. Experimental Section/Methods

Recombinant proteins were purified as previously ^[21c]^ and labelled using Oregon Green 488; details about the protein expression, purification and labeling is given in SI Section S1. Giant unilamellar vesicles of different lipid compositions were grown using the electroformation method and imaged on a Leica TCS SP8 confocal microscope; see SI Section S1 for details and for conditions of the inflation/deflation experiments in the vesicle bulk solutions. The microfluidic device as detailed in ^[26]^ was fabricated using standard soft photolithography and operated as described in SI Section S1. Fluctuation analysis was performed according to the protocol described earlier ^[41a]^, see SI Section S1. Dynamic light scattering and zeta potential measurements were performed on zetasizer Nano ZS (Malvern Instruments, Worcestershire, UK); see SI Section S1. Information about micropipette aspiration of GUVs is given in SI Section S3.

## Supporting Information

Supporting Information is available from the Wiley Online Library or from the author.

## Acknowledgements

We thank R. Seckler and S. Barbirz for kindly providing their facilities for protein purification and T. Seemann for microfabrication. We acknowledge input from R. B. Lira on the FRAP data. This work is part of the MaxSynBio consortium, which is jointly funded by the Federal Ministry of Education and Research of Germany and the Max Planck Society.

