## Supporting information for "Stepwise remodelling and subcompartment formation in individual vesicles by three ESCRT-III proteins"

### Section S1: Materials and Methods

#### *Expression and purification of recombinant proteins*

Recombinant proteins were purified as previously [1]. Briefly, proteins were induced by addition of 0.5 mM of IPTG to produce GST-rEhVps20, GST-rEhVps20t, GST-rEhVps24 and GST-rEhVps32 tagged proteins. Purified GST-tagged proteins were dialyzed against the buffer for the PreScission protease enzyme (GE-healthcare, Freiburg, Germany) and the GST-tags were removed according to manufacturer's instructions. GST-free monomers were subsequently purified by size exclusion chromatography with a Superdex 200 16/600 column (GE-healthcare, Freiburg, Germany) connected to an Äkta-Purifier FPLC (GE-healthcare, Freiburg, Germany). Proteins were stored in 50 mM Tris, 300 mM NaCl, pH = 7.4 buffer at concentrations between 60 to 10 μM.

#### *Labelling of recombinant proteins*

The recombinant proteins EhVps20, EhVps20t and EhVps32 were labelled using Oregon Green 488 (OG) (Molecular Probes-Thermo Fisher) following the manufacturer's protocol. The labelled and unlabelled proteins were separated by size exclusion chromatography with a Superdex 200 16/600 column (GE-healthcare, Freiburg, Germany) connected to an Äkta-Purifier FPLC (GE-healthcare, Freiburg, Germany). The degree of labelling was assessed according to manufacturer's instructions. In all cases, we used 1:4 ratio of labelled: unlabelled proteins to maintain activity.

#### *Preparation and imaging of giant unilamellar vesicles*

The lipids 1-palmitoyl-2oleoyl-sn-glycero-3-phosphocholine (POPC), 1-palmitoyl-2-oleoyl-sn-glycero-3-phosphocholine (POPS), cholesterol (Chol) and 1,2-dioleoyl-sn-glycero-3-phospho-(1'-myo-inositol-3'-phosphate) (PI(3)P) were purchased from Avanti Polar Lipids, Alabaster IL. In all cases, we added either Texas Red® 1,2-dihexadecanoyl- sn-glycero-3-phosphoethanolamine (TR-DHPE) (Molecular Probes) or $DiIC_{18}$ (1,1'-dioctadecyl-3,3,3',3'-tetramethylindocarbocyanine perchlorate) at a concentration of 0.1 mol% in the lipid mixtures for the visualization of the membranes. Giant unilamellar vesicles of different lipid compositions were grown using the electroformation method [2]. Briefly, 10 µl of a 4 mM lipid stock solution in chloroform were spread on indium tin oxide (ITO) coated glasses. The excess of chloroform was eliminated under vacuum at room temperature (RT) for 1h. Then, ITO-glasses were assembled with a 2 mm-thick Teflon spacer between them to form the electroformation chamber, which, if not indicated otherwise, was filled with a 600 mM sucrose solution that matched the osmolarity of the buffer containing the proteins (~650 mOsm/Kg). Osmolarities were controlled and adjusted using an osmometer (Osmomat 030, Gonotec, Germany). Finally, an electric AC-field (1.6V, 10 Hz) was applied for 1 h at different temperatures (60 °C for GUVs that contain PI(3)P and RT for the rest of the compositions). GUVs were collected and cooled to RT before use. Confocal imaging was performed on a Leica TCS SP8 confocal microscope (Mannheim, Germany). $DiIC_{18}$ was excited with a diode-pumped solid-state laser 561 nm laser, OG with a 488 nM line of Argon laser and TR-DHPE was excited with the Helium-Neon-laser at 594 nm. To avoid crosstalk between the different fluorescence signals, a sequential scanning was performed. For the $DiIC_{18}$ dye, the fluorescence signal was collected in the ranges of 580-700 nm. The fluorescence signal of OG was collected between 495-530 nm and the fluorescence signal of TR-DHPE was collected between 610-750 nm. The gain and laser intensity was maintained fixed for all experiments.

***Microfluidic chamber***

We used a microfluidic device to follow the effect of ESCRT proteins on the same GUV and to observe the action of each individual added protein. The design and fabrication of the device has been detailed elsewhere [3]. The PDMS chips were produced using standard soft photolithography as follows. Master forms at a height of 40 µm were produced on a 4” silicon wafer (Si-Mat) by UV exposure of SU8-3025 (Microchem). Before use, they were salinized in an overnight atmosphere of 1H,1H,2H,2H-perfluorodecyltrichlorosilane (ABC Chemicals) to prevent unwanted adhesion of PDMS. PDMS was mixed with the curing agent in 10:1 ratio, degassed for 30 min, and subsequently poured on top of the master form to a height of approximately 5 mm. After a further degassing for 30 min, the PDMS covered wafer was heat cured at 90 °C for 3 hr. Afterwards, the PDMS was separated from the wafer and chips were cut out. Fluidic access holes were punched with a 1.5 mm biopsy puncher (Kai Europe GmbH). A sample reservoir, made from a cut pipette tip was sealed on top of the inlet with PDMS and cured at 90 °C for 30 min. Finally, the device was assembled by bonding the PDMS chip to a glass coverslip by an air plasma treatment (Plasma Cleaner PDC-002-CE, Harrick Plasma) at 0.6 mbar for 1 min. Bonding was aided by an additional 60 °C for 2 h before use.

For the sequential protein binding experiments, microfluidic devices were coated with 2% BSA (bovine serum albumin, Sigma Aldrich) dissolved in the protein buffer (25 mM Tris, 150 mM NaCl, pH = 7.4). Then, 100 µl of GUVs, pre-deflated using a buffer of 5% higher osmolarity (adjusted with glucose), were loaded into the microfluidic device at a flow rate of 10 µl/min using a syringe pump (neMESYS, cetoni) to control the flow (Fig. S1). The GUV buffer was exchanged with 100 µl of the isotonic protein buffer (25 mM Tris, 150 mM NaCl, pH 7.4) at a flow of 5 µl/min. Afterwards, the protein EhVps20t was added to the chamber to yield a final concentration of 125 nM, with a flow of 0.1 µl/min, then EhVps32 (600 nM) and EhVps24 (200 nM) were added in that order while maintaining the flow rate in the whole experiment. Similarly, GUVs were incubated with six rounds of buffer as a negative control (Fig. S2).

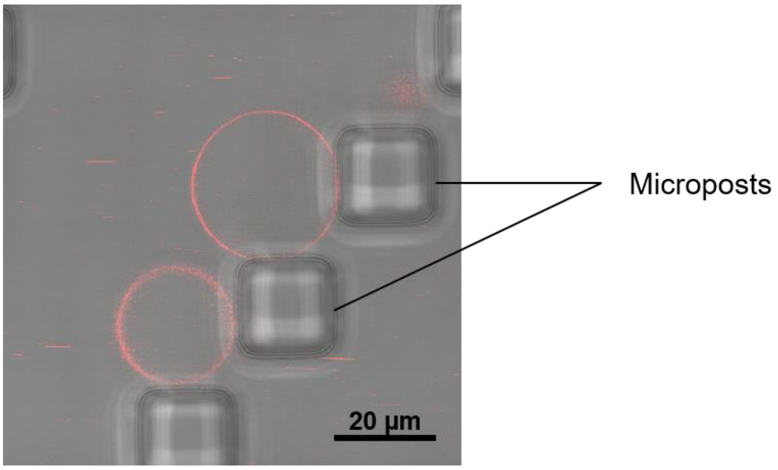


**Figure S1.** An overlay of confocal and phase-contrast image showing two vesicles trapped by the posts in a microfluidic device. The flow direction of introduced solution is from the left.

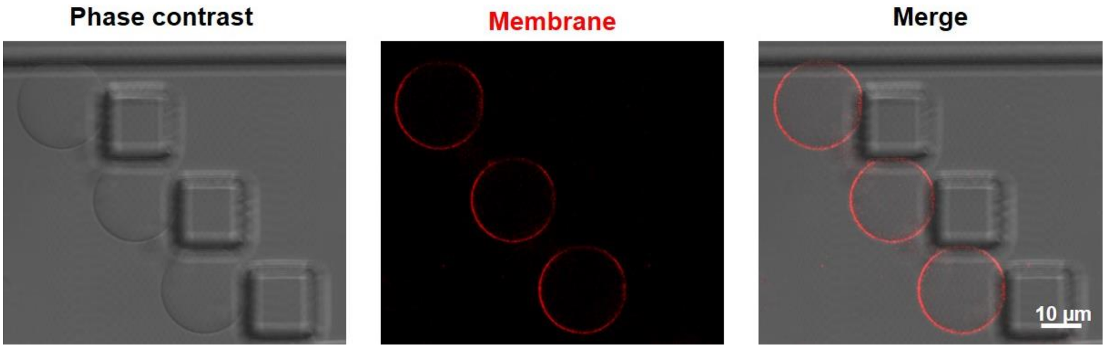


**Figure S2.** Buffer flushing and incubation of GUVs does not result in ILV formation. POPC:POPS:Chol:PI(3)P (62:10:25:3) GUVs were incubated with equivalent protein-free buffer (25 mM Tris, 150 mM NaCl, pH = 7.4) volumes as in Figure 1. Images show 3 GUVs after six rounds of buffer exchange, similar to the conditions shown in Figure 1.

***Fluctuation analysis***

Fluctuation analysis was performed according to the protocol described earlier[4]. GUVs composed of POPC:POPS:Chol:PI(3)P (62:10:25:3) and (52:10:35:3) were electroformed at 60 °C in sucrose (20 mM) and diluted in equimolar solution of glucose (containing EhVps20t at the specified concentration in the case of the lipid mixture with lower amount of cholesterol). The choice for working in low sugar concentration was set to avoid influence of gravity on the vesicle shape as well as softening effects of sugars[5]. After 5 min incubation, the vesicles were placed in a chamber made of two cover slips and a 2 mm-thick Teflon spacer, and observed under phase-contrast mode on a Zeiss Axio Observer D1 microscope using a 40× air objective. 3600 snapshots (per vesicle) were acquired with a digital camera (pco.edge, PCO AG, Kelheim, Germany) at frequency of 60 frames per second and image exposure time of 200 µs, at room temperature (~ 23 °C). The bending rigidity of vesicles in the absence and presence of EhVps20t (125 nM) was extracted. In each case, at least eight vesicles were examined. To analyse the obtained data, several normality tests were performed, namely Shapiro-Wilk, Anderson-Darling, D'Agostino-K and Chen-Shapiro and then, a two-sample test of variance was executed. Finally, a two-sample t-test was performed. All tests were accomplished at the 0.05 level (cut-off for significance).

***Dynamic light scattering and zeta potential measurements***

Large unilamellar vesicles (LUVs) were used to determine the zeta potential of the POPC:POPS (80:20) and POPC:POPS:Chol:PI(3)P (62:10:25:3) lipid mixtures. Both lipid compositions, with total lipid concentration of 0.75 mg/mL, were prepared from chloroform lipid stock solutions. The solvent was evaporated under a stream of N2 and the lipid films were further dried under vacuum for two hours to remove chloroform traces. The lipid films were hydrated in protein buffer (25 mM Tris, 150 mM NaCl, pH=7.4) to a final concentration of 0.5 mM and gently stirred for 10 min. The obtained lipid suspensions were extruded 30 times through polycarbonate membranes (Whatman, Maidstone, UK) with 100 nm pores using LiposoFast pneumatic extruder (Avestin, Otawa, Canada). The size and the zeta potential of the LUVs were measured with a zetasizer Nano ZS (Malvern Instruments, Worcestershire, UK) equipped with a 4 mW HaNe laser (632.8 nm), a detector positioned at a scattering angle of 173° and a temperature controlled cuvette holder. Three dynamic light scattering (DLS) measurements consisting of 20 runs with duration of 10 seconds at 25 °C were performed. The obtained intensity size distributions exhibited a peak at 120 nm. For the zeta potential measurements, 750 µl of the samples were loaded in DST1060 folded capillary cells with integral gold electrodes (Malvern). Three measurements each consisting of 70 runs were performed for every sample at 25 °C. Zeta potential was deduced from the electrophoretic mobility, $\mu_e$, data using the Smoluchowski and Hückel equation[6], $\zeta = 3\mu_e\eta/[2\varepsilon\varepsilon_0 f(R/\lambda_D)]$. where $\eta$ is the viscosity of the aqueous solution $\varepsilon_0$ and $\varepsilon$ are the permittivity of free space and the relative permittivity of the medium, and $f(R/\lambda_D)$ is the Henry function. The latter depends of the vesicle radius $R$ and the Debye length $\lambda_D$ of the solution. For $1 < \lambda_D < 1000$, as for the samples used here (the radius of the LUVs is 60 nm as measured with DLS), a prefactor of

$f(R/\lambda_D) = \frac{1}{6}\log(R/\lambda_D) + 1$ was taken into account in the analysis , see Ref. [7]. The Debye length $\lambda_D$ for the samples was extracted based on its measured conductivity using $\lambda_D = \sqrt{(\varepsilon\varepsilon_0 D)/K}$, where the $D$ is the diffusion constant of water $D$ = 2.299 x $10^{-9}$ $m^2/s$ [8] and $K$ is the solution conductivity which was measured $K$ = 2.87 S/m. This yields for the Debye length $\lambda_D$ = 0.75 nm. The respective zeta potentials of POPC:POPS (80:20) and POPC:POPS:Chol:PI(3)P (62:10:25:3) vesicles in the used buffer were measured to be –14 mV and –17 mV which are indistinguishable considering the instrument accuracy of ~5 mV.

***Inflation/Deflation experiments in the bulk***

POPC:POPS (80:20) GUVs were prepared by electroformation in a high osmolarity sucrose solution (~ 650 mOsm/Kg). Then, the GUVs were diluted 1:1 vol:vol in a 2 fold protein buffer (50 mM Tris pH =7.4, 300 mM NaCl, adjusted to have 5% higher osmolarity to ensure excess membrane area for bud formation) and incubated with 125 nM of EhVps20t and 600 nM of EhVps32, leaving 5 minutes of incubation between each protein addition (mixture 1). Note that incubation of the vesicles in protein-free buffer resulted in visibly fluctuating vesicles and occasional inward nanotubes with sub-optical thickness but never micron-sized buds; the vesicles were also deformed into oblates due to gravity. For the subsequent inflation/deflation steps, we used protein buffer (25 mM Tris pH =7.4, 150 mM NaCl) with the appropriate amount of proteins and adjusted the osmolarity with sucrose to achieve the work conditions indicated in the main text. This was done in order to keep the salt and protein concentrations equal throughout the whole experiment. When ILVs were visible in the GUVs, mixture 1 was diluted 1:2 (vol:vol) with a hypotonic buffer solution (~ 450 mOsm/Kg) to obtain an inflation of ~ 20% and incubated the sample for 20 min to equilibrate (mixture 2). Afterwards, the inflated GUVs of the mixture 2 were incubated 1:1 with a hyperosmotic buffer (~ 630 mOsm/Kg) to reach a deflation of ~10%.

***Fluorescence recovery after photobleaching (FRAP)***

FRAP measurements were performed on intraluminal buds as well as on the GUV surface. Movement of the vesicles in the sample because of convection hindered the experiments. To minimize the GUV movement during the measurements the vesicles were immobilized in agarose following the methodology in Ref. [9]. Briefly, electroformed vesicles (grown in sucrose) were incubated with 125 nM EhVps20t and 300 nM OG-EhVps32 for 10 min. 8 µL of the GUV-protein mixture were deposited on a cover glass (passivated with BSA) and immediately 2 µL 0.5 % w/v preheated agarose solution diluted in the protein buffer were added. The solutions inside and outside the GUVs were osmotically balanced. Confocal microscopy images were recorded at 1000 Hz with a pinhole size of 1 Airy unit in bidirectional mode and with an image size of 512 x 512 pixels and at room temperature (23 °C). OG-EhVps32 was exited using the 488 nm line of the argon laser and the fluorescence signal was detected in the range 493-550 nm, while $DiIC_{18}$ was excited with the 561 nm laser line, and the fluorescence signal collected at 567-628 nm. Pre-bleaching, 10 frames at attenuated laser intensity (below 5%) were recorded. The photobleaching was performed for 505 ms (3 frames) at 100% laser intensity using a circular region of interest (ROI) of nominal radius $r_n$ = 3.6 µm, and 3 µm in the case of intraluminal buds or the top of the GUVs, respectively. The post-bleach recovery images were then recorded at the initial attenuated laser intensity for 100 frames. The diffusion coefficient, $D$, from the FRAP recovery performed on the top of the GUV was extracted using $D = r_e^2 + r_n^2/8t_{1/2}$ were $r_e$ and $r_n$ are the effective and the nominal radii, respectively and $t_{1/2}$ is the halftime for the dye photorecovery [10]. Some of the bleached buds (roughly 70%) showed non-monotonous recovery curves, because the buds were diffusing and leaving the focal plane during the measurement; this could be also observed in the images acquired under phase-contrast imaging. Such recovery data were discarded to avoid contribution to the signal due to bud defocusing.

**Section S2. Measuring the protein coverage at the vesicle membrane.**

GUVs composed by 80 mol% POPC, 20 mol% POPS and different concentrations of OG-DHPE (0.1, 0.25, 0.5 and 0.75 mol%) were generated by electroformation as detailed before. GUVs were observed with a Leica TCS SP5 confocal microscope. In order to calculate the absorption of EhVps20t at the membrane, a calibration curve of the dye was generated. Following the procedure from Weinberger *et al.*, [11], we first checked that the intensity of the Oregon green 488-labelled protein (OG-EhVps20t) behaves linearly with the concentration of the fluorophores present in the sample. Thus, we measured the fluorescence intensity of different concentrations of the labelled protein using confocal microscopy while maintaining the same microscope settings and objective for the whole quantifications (see Section S1). Figure S3 shows the linear dependence of the intensity obtained from these experiments.

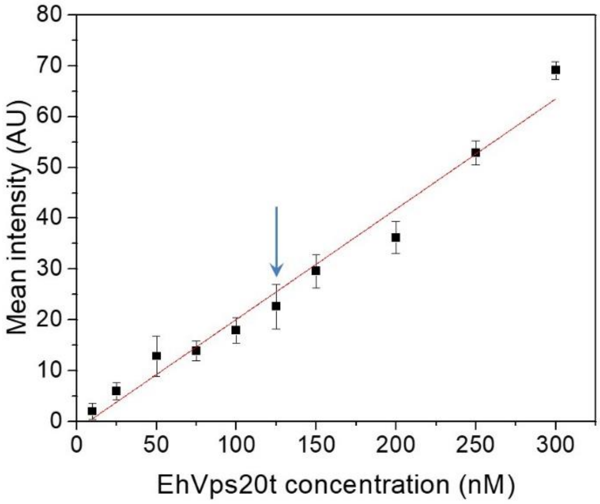


**Figure S3.** Linear dependence of the bulk fluorescence intensity of OG-EhVps20t mixed with unlabelled EhVps20t (1:4, v:v) dissolved in 25 mM Tris, 150 mM NaCl, pH = 7.4 at room temperature. Arrow points to the typical working concentration. The red line is a linear fit. The error bars represent the standard deviations.

Next, for measuring the intensity at the level of the membrane, we used the “Radial Profile Extended Plugin” from Philippe Carl available in the ImageJ homepage. We prepared GUVs made of POPC:POPS (80:20) including various concentrations of the labelled lipid OG-DHPE (0.01, 0.025, 0.05 and 0.075 mol%) in the mixture. The radial intensity profile of the different GUVs was obtained at the equatorial plane. To avoid polarization issues resulting from the dye orientation with respect to the membrane, a complete circle was centred at the vesicle centre and intensity radially averaged. An example of an intensity profile is shown in Fig. S4. The integrated peak area is then taken as the intensity for the specific fluorophore concentration in the membrane. The scatter in the data obtained in Fig. 2a in the main text results from the fact that we have analysed vesicles of different sizes whereby out-of-focus fluorescence contributions vary. To use the obtained calibration curve in Fig. 2a and deduce the amount of bound protein, we have divided this intensity value by a factor of 2 as to correspond to fluorescence from the outer vesicle leaflet (as is the case of adsorbing OG-labelled protein).

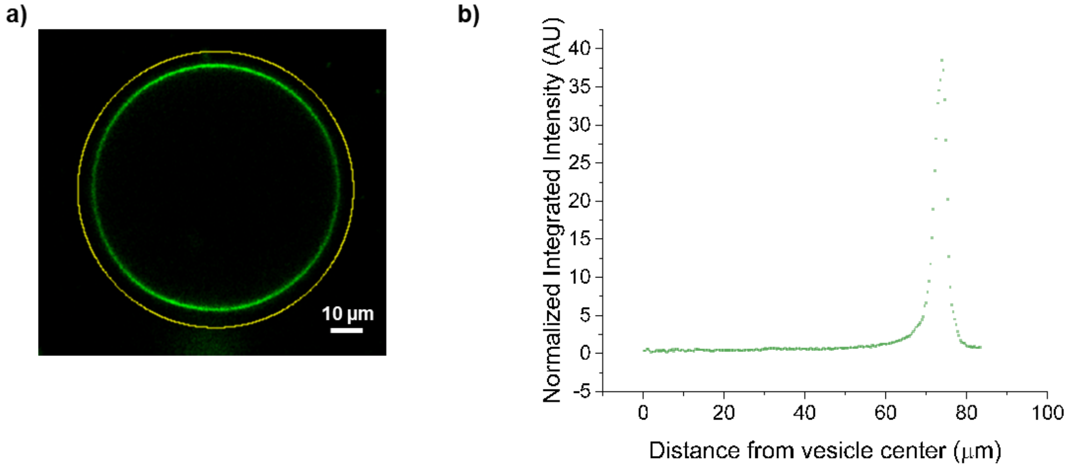


**Figure S4.** (a) Representative confocal image of a vesicle labelled with OG-DHPE used for the standard curve generation; the polarization effect is visible from the angular dependence of the florescence. (b) Typical intensity radial profile. The value of intensity was obtained by measurement of the area under the curve generated in the plot.

To calculate the adsorption of OG-EhVps20t at the membrane of the GUVs, we took into consideration the signal from the unbound protein in the bulk solution and subtracted it. Figure S5 illustrates the procedure that we followed. The protein coverage on vesicles prepared from POPC:POPS:Chol:PI(3)P (62:10:25:3) was found to be similar as expected from the comparable surface charge, see Fig. S6.

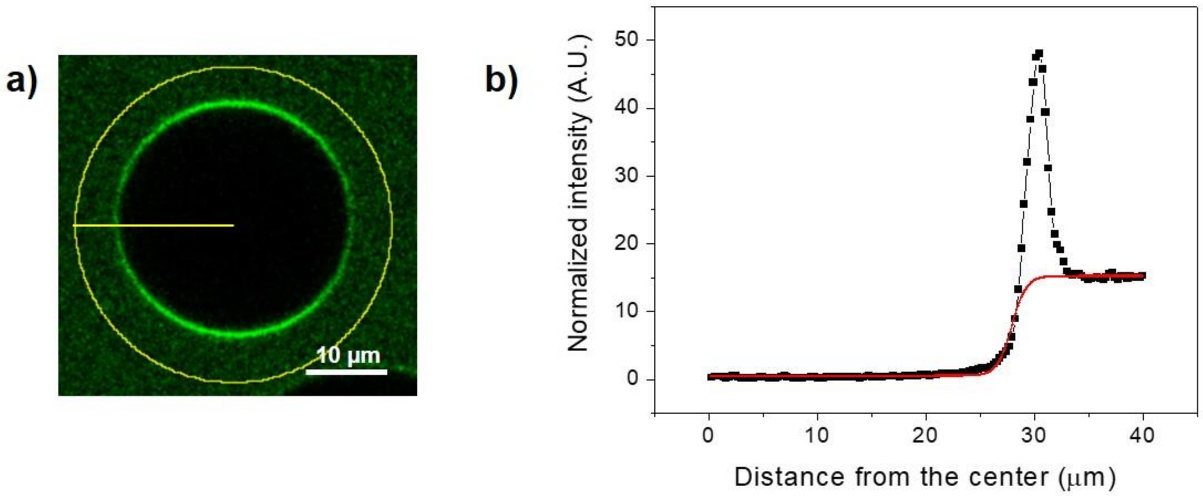


**Figure S5.** (a) Typical confocal image of a GUV used for measuring the coverage concentration of OG-EhVps20t at the membrane. (b) Radial intensity profile (black squares) and subtracted intensity of the non-adsorbed protein in the bulk (red curve).

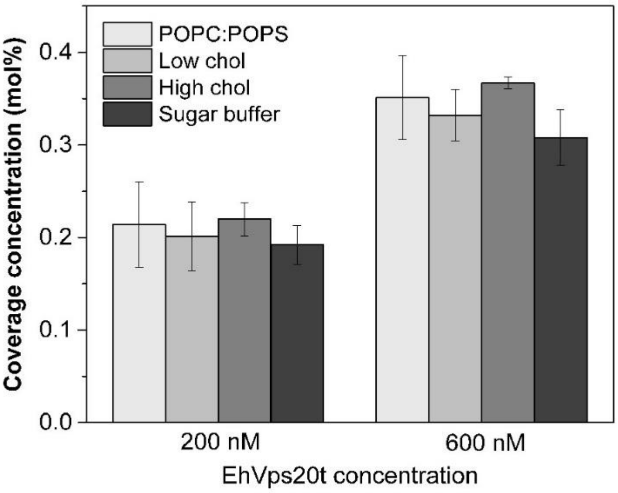


**Figure S6.** Protein coverage of EhVps20t at two different concentrations of the protein present in the bulk. In order from left to right, different types of conditions were tested (1) POPC:POPS (80:20) GUVs, (2) POPC:POPS:Chol:PI(3)P (62:10:25:3) GUVs (Low chol) and (3) POPC:POPS:Chol:PI(3)P (6:10:35:3) GUVs (High chol), incubated in the normal protein buffer (25 mM Tris, 150 mM NaCl, pH = 7.4). The darker bar, represent results from POPC:POPS:Chol:PI(3)P (62:10:25:3) GUVs incubated in sugar buffer (600 mM Sucrose). At least twenty vesicles were measured for each condition, standard errors are shown for each bar.

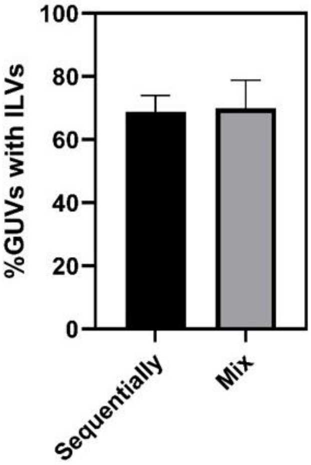


**Figure S7.** Effect on ILV formation by protein addition approach – sequentially or premixed**.** POPC:POPS:Chol:PI(3)P (62:10:25:3) GUVs were incubated with 125 nM of EhVps20t, 600 nM of EhVps32 and 300 nM EhVps24, either sequentially leaving 5 min of incubation between each protein addition or together in a mix leaving 15 min incubation at the end. Data represent the mean and standard error of three independent experiments where 30 GUVs of each replicate were screened.

**Section S3. Micropipette aspiration of GUVs.**

Micropipettes were pulled from borosilicate capillaries (1B100-4, World Precision Instruments Inc.) with a pipette puller (Sutter Instruments, Novato, CA) and then shaped with a microforge (Narishige, Tokyo, Japan). The inner pipette diameter varied in the range of 10–15 µm. The pipette tips were incubated in protein buffer (25 mM Tris, 150 mM NaCl, pH=7.4), containing 1% BSA to prevent adhesion of the vesicle membrane to the pipette. After the incubation, the micropipette was rinsed with the protein buffer to remove the free BSA. Aspiration of GUVs was performed in a homemade experimental chamber with volume of 0.5 ml. The chamber was built from two parallel glass coverslips separated by a Teflon spacer with opening from one side for inserting the micropipette. To prevent vesicle adhesion, the glasses were passivated with 1% BSA solution prior to chamber assembly. The vesicles were observed on a Leica TCS SP5 confocal microscope (Mannheim, Germany), equipped with 40 × objective. $DiIC_{18}$ was excited with a diode-pumped solid-state laser 561 nm laser and the fluorescence signal was collected in the ranges of 580-700 nm. The micropipette was operated using a three-dimensional micromanipulator system (Sutter Instruments, Novato, CA) mounted on the microscope. The aspiration pressure in the micropipette was controlled by changing the height of a water reservoir mounted on a linear translational stage (M-531.PD; Physik Instrumente Germany). Equilibrium height of the water reservoir corresponding to zero pressure across the pipette tip was set prior to each measurement.

The membrane tension was assessed as

$$\Sigma = \frac{\Delta P\ R_p}{2(1 - R_p/R_{ve})}$$

where $\Delta P$ is the suction pressure, $R_{ve}$ and $R_p$ are respectively the radii of the spherical vesicle and the micropipette. Since the ILVs are connected via a narrow neck to the mother vesicle the upper limit of the spontaneous curvature $m$ can be estimated from the neck closure condition $m \leqslant M_{ne} + \frac{1}{2R_W} = \frac{1}{2}\ (1/R_{ve} - 1/\overline{R_{ILVs}})\ + \frac{1}{2R_W}$, where $\overline{R_{ILVs}}$ is the average radius of all ILVs and $R_W = (2\kappa/|W|)^{1/2}$ is the adhesion length with the adhesion strength $|W|$ between the membrane and the protein assembly[12]. The estimates for the spontaneous curvature $m$ in the main text have been obtained by omitting the term $\frac{1}{2R_W}$ reflecting our ignorance about the adhesion strength $|W|$ The spontaneous tension is then $\sigma_m = 2\kappa m^2$, where $\kappa$ is the bending rigidity of the membrane. This estimate is valid for homogeneous membranes (same $\kappa$ and $m$ for vesicle and buds). The area expansion ($\Delta A/A_0$) was calculated using equation 46 from [13].

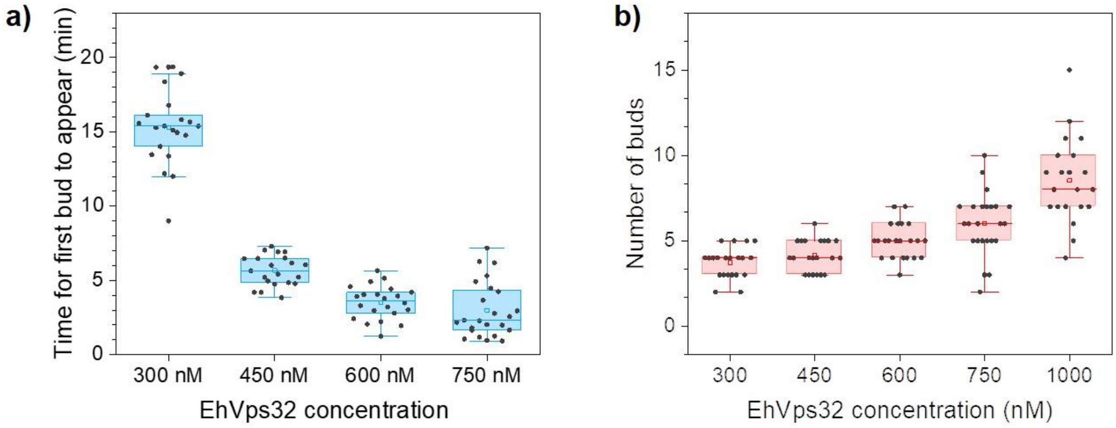


**Figure S8.** Kinetics of bud formation regulated by EhVps32 concentration. POPC:POPS:Chol:PI(3)P (62:10:25:3) GUVs were incubated with 125 nM of EhVps20t for 5 min followed by different concentrations of EhVps32 as shown in the x-axis. At least 10 vesicles of three independent preparations were followed over time and (a) the time that took for the first bud to be observed is plotted. At higher concentrations of EhVps32, namely 1000 nM, the process was faster than 1 minute and the exact time frames could not be determined because of difficulties in experimental handling. (b) Total number of buds detected in each GUV after 30 min of incubation with EhVps32. Black data points show the single measurements for each condition. Boxes show the range from 25-75% of data and the median line.

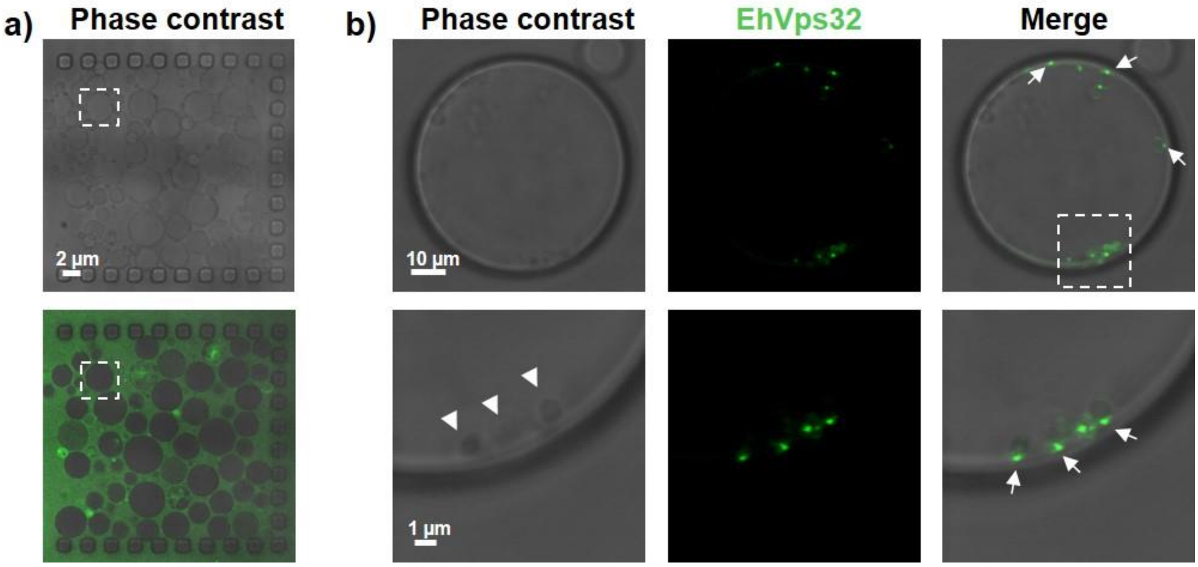


**Figure S9.** EhVps32–enriched domain formation in GUVs. (a) POPC:POPS:Chol:PI(3)P (62:10:25:3) GUVs were loaded on a microfluidic chamber at a flow rate of 10 µl/min. Then 125 nM of EhVps20t and 300 nM of EhVps32 (20% of EhVps32 was labelled) were subsequently flushed at a constant rate of 0.1 µl/min. (b) Close-up views of the vesicle in the dashed region in (a) after the excess of proteins was washed away, showing protein-domain formation (arrows) in the site where buds (arrowheads in phase-contrast image) are generated.

```
ScVps20     1 MGQKS----S-----------KVHITKTDRAILEVKRS-KDEIHKFTRRTDNLILVEKSQ
CHMP6       1 MGNLFGR-KK-----------QSRVTEQDKAILQLKQQ-RDKLRQYQKRIAQQLERERAL
EhVps20     1 MLNRFIGKNEEPSIDETHAILESRVTDLDQKIVDLDRQIRQNIEMSHKTTGAESQ-----

ScVps20    45 LKDLIRKNPENYKSNMKVRFLLKRIHYQEHLLQQASDQLINLENMVSTLEFKMVEKQFIN
CHMP6      48 ARQLLRD-----GRKERAKLLLKKKRYQEQLLDRTENQISSLEAMVQSIEFTQIEMKVME
EhVps20    56 ------------RYKQKALMLLKQKKQHEQHRDITERHQMNLENVQFQVDQMKDYKKVYD

ScVps20   105 GLKNGNEILKKLNKEFS--NVDELMDDVQDQIAYQNEINETLSRSLVGTSNYEDDLDKEL
CHMP6     103 GLQFGNECLNKMHQVMSIEEVERILDETQEAVEYQRQIDELLAGSFT--QEDEDAILEEL
EhVps20   104 AMAASNRQMKLISRQFDINSIEDVQDSMMNFAEDQQEFGDILSQDIT--NMDLNELDDEL

ScVps20   163 DALESELNPEKMNNAKVANMPSTEGLP-SLPQGEQ-TEQKEREEFATEERSDTKEPLALL
CHMP6     161 SAIT----QEQIELPEVPSEPLPEKIPENVPVKAR-PRQ--------------AE-LVAA
EhVps20   162 AELG----DSMFE-GDI-QIPESDGFG-SVPITNEPFRQKEE------SQPDDKK-FA--

ScVps20   221 S
CHMP6     201 S
EhVps20   206 I
```

**Figure S10.** EhVps20 lacks the glycine residue in its N-terminal necessary for protein myristoylation. Multiple sequence alignment for Vps20-homologues in yeast (ScVps20), human (CHMP6) and *E. histolytica* (EhVps20). Glycine residues present in EhVps20 are displayed in red. Arrow indicates the N-terminal glycine necessary for protein myristoylation. Sequence alignments were performed with Clustal Omega and edited in the BoxShade server (https://embnet.vital-it.ch/software/BOX_form.html).

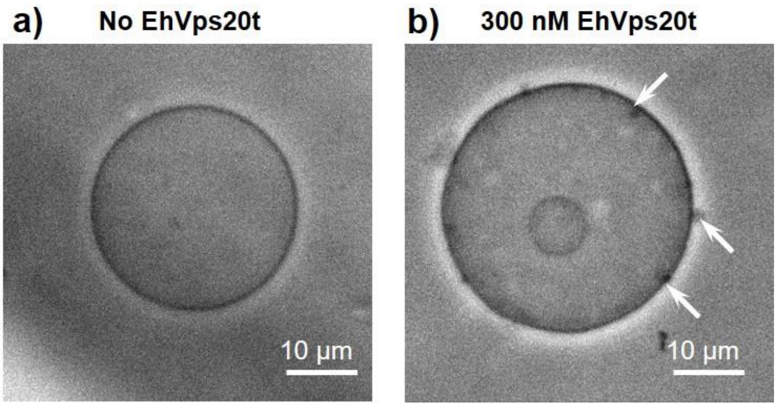


**Figure S11**. EhVps20t clusters in the membrane of GUVs at high protein concentrations. Electroformed POPC:POPS:Chol:PI(3)P (62:10:25:3) GUVs incubated in (a) a isoosmolar sucrose buffer (no EhVps20t) or (b) 300 nM of EhVps20t, observed under phase contrast. Arrows show protein clusters at the membrane.

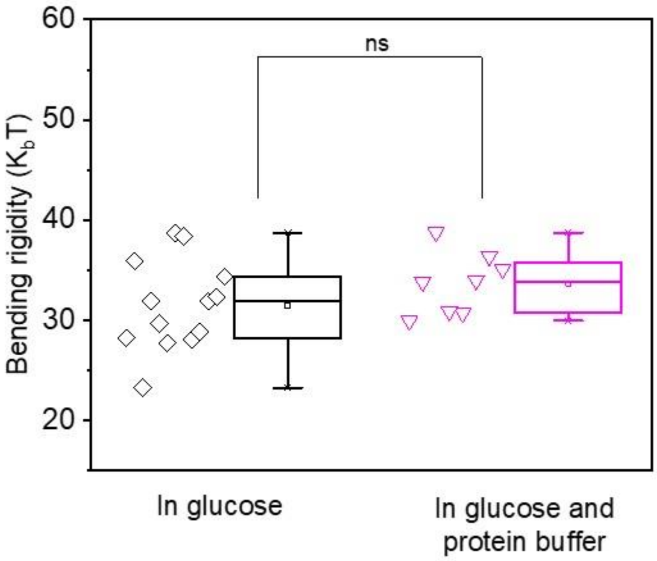


**Figure S12.** Bending rigidity of protein-free membranes. The vesicles were formed of POPC:POPS:Chol:PI(3)P (62:10:25:3) in 20 mOsm/Kg sucrose solution. The GUVs for the data set indicated by the black diamonds (the same data as "Low Chol" in Fig. 5c in the main text) were diluted in isotonic glucose solution, while the vesicles used for the data in magenta triangles were diluted in a solution containing protein buffer and glucose. The latter solution represents the scenario when 125 nm EhVps20t was used (see Fig. 5c in the main text).

### Supplementary Movies

**Movie S1.** The video shows the complete scan of an electroformed POPC:POPS:Chol:PI(3)P (62:10:25:3) GUV labelled with 0.1 mol% TR-DHPE after the consecutive incubation with 125 nM of EhVps20t (20% of the protein was labelled with Oregon green; merged signal is shown), 600 nM EhVps32 and a washing step between the incubation of the two proteins as done in Fig. 1. The intraluminal buds are still attached to the GUV membrane and are seen in different planes of the vesicle. They move and occasionally appear elongated because of the present flow and the slow scanning speed. The GUV diameter is approximately 30 µm.

**Movie S2.** The video shows ILVs movement inside a GUV trapped in a microfluidic device monitored under phase contrast microscopy. The small light and dark spots which are observed to appear/disappear and move inside the trapped GUV represent ILVs that come in and go out of focus. The vesicle is of 20 µm in radius. The actual duration of the recorded sequence in the movie is 20 s. During the recording, another vesicle is pressed against the main one by the flow but does not adhere to it as seen in the end of the movie, when it is flushed away.

**Movie S3.** The video shows the complete scan of an electroformed POPC:POPS:Chol:PI(3)P (62:10:25:3) GUV after the deflation step as in the bulk experiments of Fig. 3 (same vesicle as in Fig. 3a, bottom panel). The GUV was in protein buffer (25 mM Tris, 150 mM NaCl, pH 7.4) with osmolarity adjusted to 822 mOsm/Kg using glucose. The vesicle is 15 µm in radius.

**Movie S4.** The video shows three complete scans of the GUV from Fig. 3b in the main text before inflation (top left), after inflation (bottom left) and after deflation (top right).
